# Representation Transfer via Invariant Input-driven Neural Manifolds for Brain-inspired Computations

**DOI:** 10.1101/2025.07.27.666997

**Authors:** Tie Xu, Shengdun Wu, Junwen Luo, Feng Lin, Junjie Jiang, Guozhang Chen, Yao Yang, Jieping Ye, Dongping Yang

## Abstract

Domain adaptation is a core challenge for embodied AI deployed in unpredictable, noisy environments. Conventional deep models degrade under domain shifts and require costly retraining. Inspired by biological brains, we propose a modular framework where each module is a recurrent neural network pretrained via a simple, task-agnostic protocol to learn robust, transferable features. This shapes stable yet flexible representations as invariant input-driven continuous attractor manifolds embedded in high-dimensional latent space, supporting robust transfer and resilience to noise. At deployment, only a lightweight adapter needs training, enabling rapid few-shot adaptation. Evaluated on the DVS Gesture benchmark and a custom RGB rehabilitation dataset, our framework matches or surpasses leading C3D and ViViT models while using ten times fewer parameters and only one training epoch. By unifying biologically inspired attractor dynamics with cortical-like modular composition, our approach offers a practical path toward robust, continual adaptation for real-world embodied AI.

**Teaser:** Invariant input-driven attractors enable representation transfer and domain adaptation in brain-inspired AI.

## 1 Introduction

Artificial Intelligence (AI) systems deployed in embodied contexts–such as robotics, autonomous vehicles, and wearable assistants–must operate reliably in open-ended, unpredictable environments. In such settings, models face pervasive challenges, including sensor noise, adversarial perturbations, data imbalance, and environmental non-stationarity (*1*). Achieving robust performance under these conditions demands rapid domain adaptation (*2*), especially when training data are scarce or the environment shifts over time.

However, most modern AI systems rely on end-to-end training, assuming identical training and test distributions (*2*). This static paradigm severely constrains adaptability, making conventional task-specific architectures brittle under distributional shifts (*3*). Reservoir computing (*4*) offers a promising alternative for fast adaptation in temporal tasks: by projecting inputs into a rich, fixed recurrent “reservoir” and training only a lightweight readout, it dramatically reduces training time and data requirements (see Supplementary Note 1 for details). Yet, reservoirs with random internal weights often lack the structured representations and feature learning needed for solving more complex, transferable, or generalizable tasks.

In contrast, biological nervous systems exhibit remarkable adaptability and generalization, even with limited supervision (*5*). This flexibility stems from evolutionarily conserved neural circuits (*6*) that encode richly structured dynamics, frequently organized as attractors–low-dimensional, stable patterns of neural activity (*7*). Attractors have been proposed as reusable cognitive “symbols” for fundamental concepts or operations (*8, 9*), enabling circuits to be flexibly recombined across tasks rather than rebuilt each time. Their inherent noise tolerance and representational stability make them well-suited for robust, modular computation and for transferring representations to new tasks and contexts (*10,11*). However, canonical attractor models are typically designed with slow autonomous convergence (*12, 13*), while real-world sensory tasks demand rapid, input-driven responses (*7, 13*). This timescale mismatch leaves most current AI architectures without mechanisms to reach stable yet adaptive dynamics (*12*).

To harness the generalization power of attractor dynamics, we propose a brain-inspired framework that explicitly embeds input-driven attractor manifolds (*12*) as modular, transferable representations for robust domain adaptation. Drawing on insights from neuroscience and dynamical systems, we design recurrent neural network (RNN) modules pretrained on simple synthetic sequences that capture core spatial and temporal symmetries, rather than on task-specific labels. This task-agnostic pretraining shapes each module’s high-dimensional state space into structured, low-dimensional manifolds–rings, cylinders, and tori–that represent interpretable motion primitives such as direction, velocity, and spatial position. Crucially, these input-driven “slow manifolds” (*14, 15*) support smooth, stable dynamics that remain responsive to rapidly changing inputs, allowing the network to robustly encode continuous task variables and adapt to shifting conditions. Evaluations on DVS gesture and RGB rehabilitation action-recognition tasks show that these manifolds preserve their structure–anchored to the same interpretable motion variables–and maintain strong performance even under severe perturbations such as dropped or corrupted frames.

The resulting attractor modules can be composed flexibly within a modular architecture, with each specializing in distinct spatiotemporal features. A Hebbian-inspired fast-adaptation mechanism enables lightweight task reconfiguration without disrupting the stable pretrained dynamics. This design achieves strong few-shot performance–matching or exceeding conventional deep models– while using an order of magnitude fewer parameters and requiring only minimal additional training. By embedding input-driven attractor dynamics within a brain-inspired modular architecture, our framework enables systems that are inherently robust, interpretable, data-efficient, and energy-efficient–qualities crucial for edge computing and continual learning. Therefore, our results suggest a new design paradigm for embodied AI: one that grounds adaptive computation in the dynamical principles of biological brains.

## 2 Results

### 2.1 Framework overview

Our approach is motivated by the observation that many human actions–particularly in everyday gestures or rehabilitation contexts–are characterized by constrained, low-complexity movements that closely resemble rigid-body dynamics (*16, 17*). According to classic kinematic theory (*17*), such movements can be decomposed into basic primitives: *translation, rotation*, and *oscillation*. This suggests that recognizing these simple actions can be reduced to identifying a few core spatiotemporal observables, such as spatial location and configuration of body parts, and key temporal features such as direction and velocity of movement (Fig. 1a).

**Figure 1:**
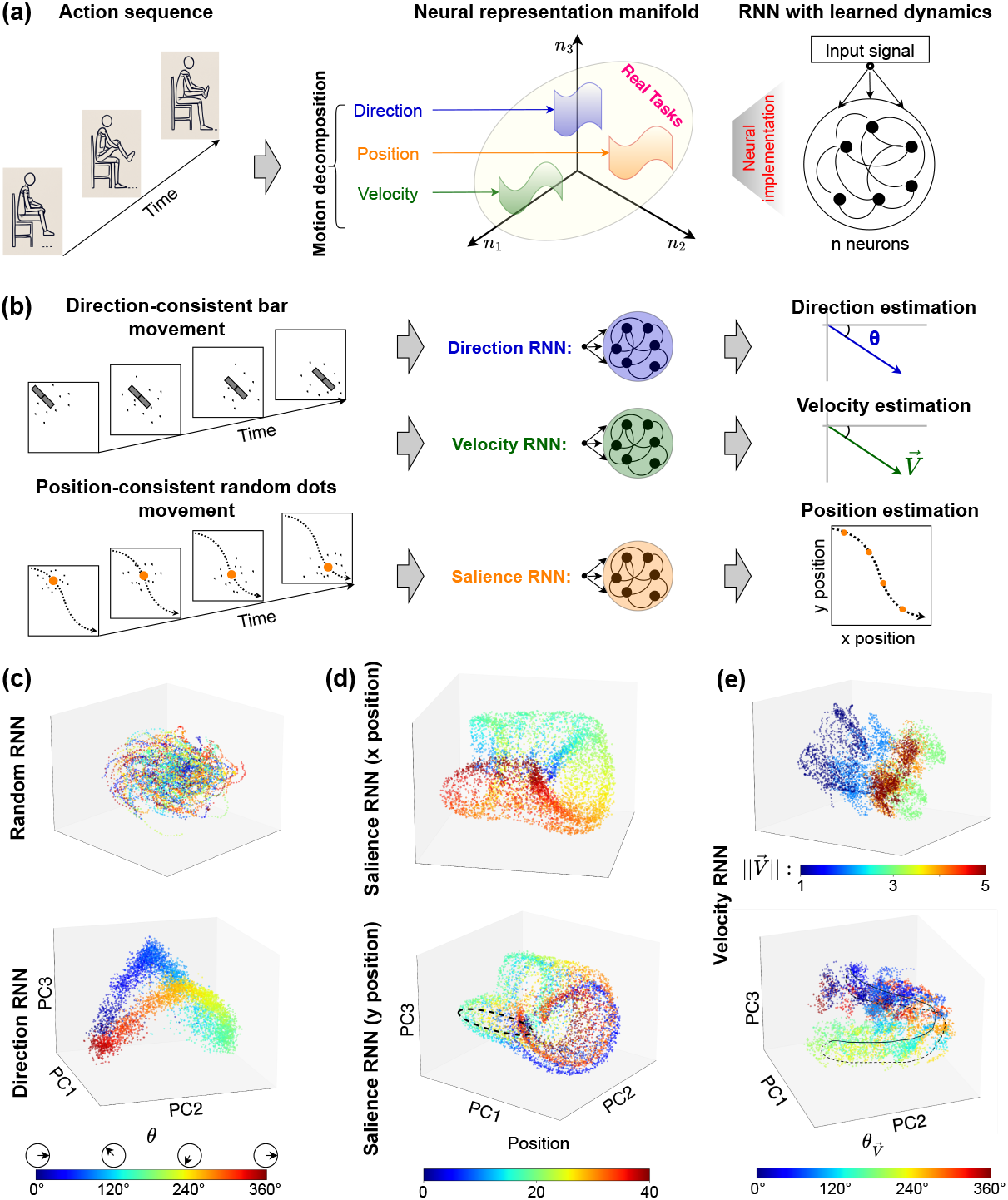
Overall concept, pretraining schema, and emergent neural manifolds. (**a**) Conceptual overview: Action sequences are decomposed into core motion attributes–direction, velocity, and position–captured by specialized, pretrained RNN modules as low-dimensional neural manifolds. Pretraining tasks: The direction RNN learns to estimate motion direction *θ* from bar motion; The velocity RNN is trained to estimate motion velocity, including motion speed 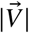 and velocity direction 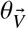. The salience RNN tracks the center position (*x, y*) from moving dot patterns. (**c**-**e**) Principal component (PC) projections of the dynamics in RNNs: (**c**) Direction RNN: The trained RNN (bottom) forms a smooth ring manifold aligned with input direction, unlike the unstructured dynamics of a random RNN (top); (**d**) Salience RNN: Smooth toroidal manifold aligned with *x* (top) and *y* (bottom) positions; Black dashed line indicates trajectory at a fixed azimuthal angle through the toroidal manifold. (**e**) Velocity RNN: joint and structured manifold encoding of speed 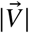 (top) and velocity direction 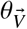 (bottom).

We hypothesize that neural circuits pretrained to extract these elementary features from synthetic inputs can develop internal dynamics that serve as reusable inductive biases, enabling generalization to more complex, naturalistic settings with more robustness. To test this, we pretrain each vanilla RNN on synthetic, task-agnostic video sequences designed to isolate a specific motion attribute (*18*). Inspired by principles of developmental learning in the visual system (*19, 20*), this pretraining process is designed to shape each RNN into a domain-specialized encoder for one of three fundamental features: motion *direction, velocity*, and *spatial salience* (Fig. 1b). This biologically inspired pretraining paradigm sculpts smooth, structured, low-dimensional dynamics within each RNN, yielding stable, interpretable manifolds aligned with physical motion parameters. Importantly, these representations emerge independently of any specific downstream task, allowing each module to function as a general-purpose, transferable component for robust, modular composition. Together, the pretraining and flexible late-on integration make up our Pretrained Reservoir Group (PRG) framework (see Sec. 2.5 for details). At deployment, the pretrained modules are assembled into a task-specific pipeline for action recognition, incorporating mechanisms for input alignment, feature encoding, and Hebbian-inspired adaptive decoding. The resulting system supports strong generalization with minimal supervision, even under distributional shifts or limited data.

### 2.2 Hidden representations

Each module was initialized with random weights and pretrained with correspondent binary videos to extract a specific motion-related feature (see Sec. 5 for details):

- **Direction RNN**: Trained on *T* = 100-frame clips of a bar moving in one of 12 evenly spaced directions, with direction held constant for 20-frame intervals. Noise was introduced via additive Bernoulli flips (probability *p* = 0.01). The RNN predicted the direction *θ* at each timestep, binned into 12 classes.
- **Velocity RNN**: Trained on sequences where a bar moved at one of five discrete speeds and 12 directions. The RNN predicted both speed and direction at each timepoint, achieving ∼ 88% accuracy on held-out sequences.
- **Salience RNN**: Trained on dot-cloud clips with a smoothly moving Gaussian centroid. The RNN estimated the centroid’s (*x, y*) position on a discretized 2D grid, reaching ∼ 86% accuracy on test data.

This pretraining paradigm addresses challenges associated with task-specific end-to-end training– such as vanishing or exploding gradients (*21*)–and instead induces biologically inspired dynamics aligned with kinematic primitives. Among the attributes, direction selectivity is particularly crucial, consistent with its prevalence in early visual processing across species (*16, 22–24*).

To visualize the emergent dynamics from a randomly initialized state (Fig. 1c, top), we recorded the hidden states of each pretrained RNN and projected them into low-dimensional subspaces using principal component (PC) analysis (Fig. 1c-e). Pretraining led to striking structural transformations in the state space: 1) In the **Direction RNN**, the hidden states formed a smooth ring-like manifold, where each position encodes a distinct motion angle (Fig. 1c, bottom). This ring attractor mirrors those observed in biological circuits encoding head direction or angular velocity (*7*). 2) In the **Salience RNN**, the hidden states formed a toroidal manifold in which different positions along the surface map to (*x, y*) spatial locations, effectively embedding 2D positional information into a continuous, low-dimensional latent structure (Fig. 1d). 3) The **Velocity RNN** exhibited a cylindrical manifold, with angular components encoding motion direction and radial or axial coordinates representing speed (Fig. 1e).

In all cases, the relevant motion attribute served as an *order parameter*, around which neural activity self-organized into a smooth, low-dimensional attractor manifold. Thus, pretraining on synthetic, attribute-specific tasks can reliably sculpt modular RNNs into interpretable dynamics.

### 2.3 Probing hidden dynamics

We now probe internal dynamics of each RNN by examining their properties under internal state perturbation and input shift perturbation (Fig. 2a-b; Sec. 5). Across all modules, we find that emergent manifolds are low-dimensionally structured, input-anchored, and resilient to both internal noise and external variation. They exhibit fully *input-driven* dynamics, where the internal state is tightly coupled to the current stimulus rather than maintained autonomously. For instance, in the **Direction RNN** (Fig. 2a), presenting a directional input (e.g., *I*_*θ*_ = 90^°^) reliably positions the network’s state on a specific location along the ring manifold. Removing the input causes the activity to rapidly decay to baseline, confirming that the representation is anchored to external stimuli, rather than self-sustained. It contrasts with traditional autonomous attractors, and can be recognized as input-driven continuous attractor dynamics for encoding rapidly changing streams.

**Figure 2:**
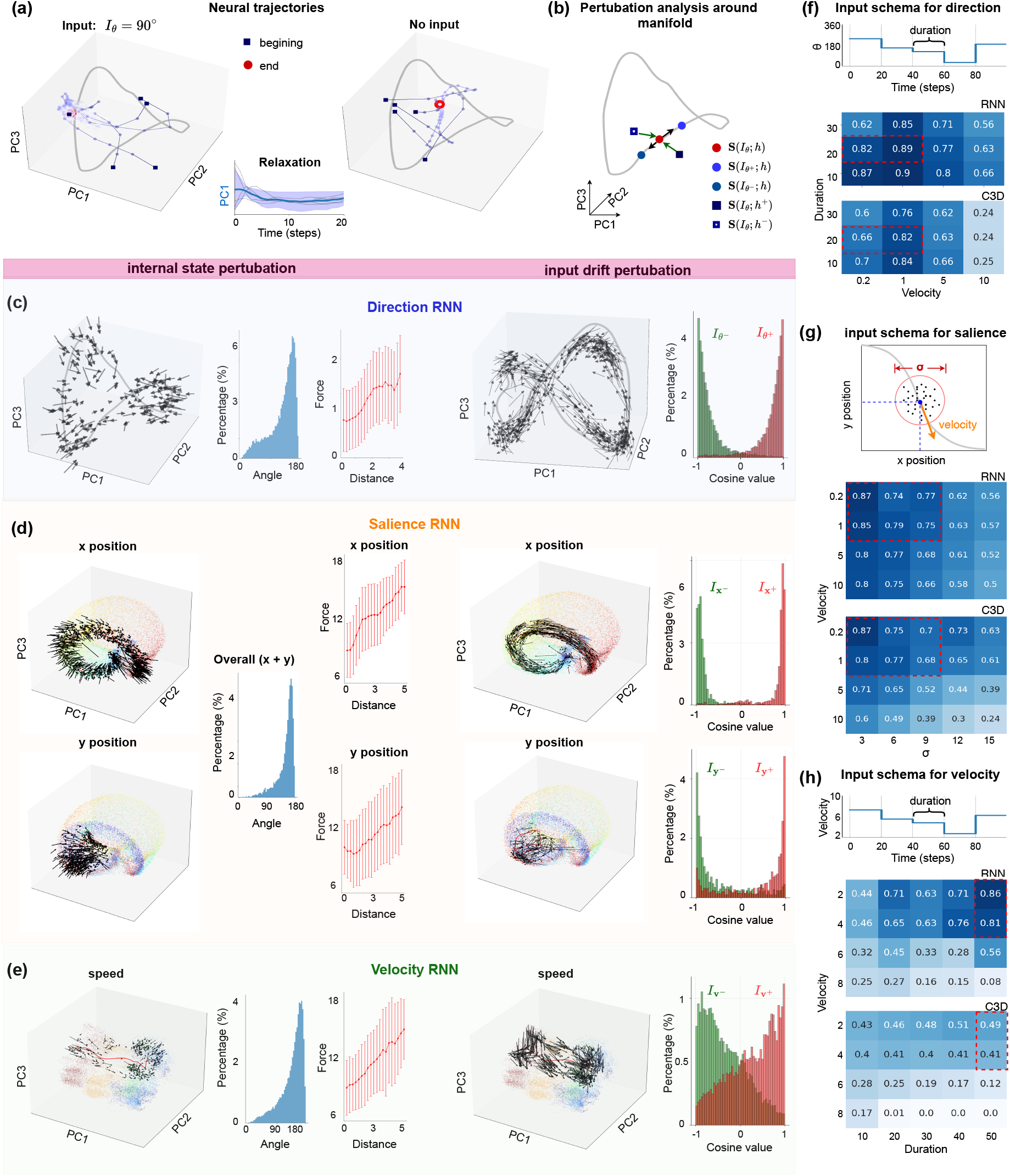
Neural dynamics around attractor manifolds in direction (**c**), salience (**d**), and velocity (**e**) RNNs. (**a**) Input-driven dynamics: activity converges to a stable point on the ring manifold under a fixed direction (*I*_*θ*_ = 90^°^) and decays when input is removed. (**b**) Perturbation types: internal (off-manifold) and input (along-manifold) shifts. (**c**) Perturbation analysis for direction RNN: Internal stability analysis including vector fields visualization, force-distance profiles, and angle histograms show recovery to the manifold, using slow manifold analysis (see Methods). Input shift analysis including vector fields visualization, and statistics of vector field relative to the tangent direction of the manifold with two opposite input offsets. (**d**) Same analyses for the salience RNN; (**e**) for the velocity RNN. In (**d**-**e**), point colors in PC spaces match Fig. 1. (**f**-**h**) Generalization accuracy across input direction, duration, and velocity variations with reference to C3D network trained in the same paradigm. Training conditions are red squared.

These input-driven dynamics confer two key advantages: robustness against internal noise and responsiveness to input shifts (Fig. 2b). Perturbation analyses reveal two core dynamical properties (Fig. 2b-e): 1) **Stability to internal perturbations**: When the hidden state is perturbed away from the attractor manifold under fixed input, network activity rapidly contracts back within 5 ∼ 10 time steps, converging toward the input-anchored attractor region on the manifold (Fig. 2c–e, left PC panels). Vector field visualizations confirm this strong restorative behavior toward attractor regions aligned with the encoded attributes (e.g., direction, location, or speed): vectors point inward toward the attractor, with angles concentrated near 180^°^ relative to the outward normal direction (angle distributions in Fig. 2c-e). Moreover, the restoring force increases proportionally with distance from the manifold (Force vs. Distance plots in Fig. 2c-e). Once on the manifold, neural trajectories settle near the attractor and exhibit slow tangential evolution–hallmarks of continuous attractor dynamics. **2) Adaptability to input drift**: When external inputs slightly shift (e.g., motion angle shifts by ±30^°^, spatial position by ±1 unit, or velocity level by ±1), neural trajectories respond with smooth, tangential flow along the manifold’s surface (Fig. 2c-e, right PC panels). This is evidenced by high cosine similarity between the flow direction and the local tangent of the manifold, indicating that the encoding geometry is preserved. Such tangential adaptation allows the network to continuously track and interpolate dynamic stimuli without losing representational stability.

These shared dynamical properties yield strong generalization even under test conditions far beyond the pretraining ones (Fig. 2f-h): 1) The **Direction RNN** maintains over 80% decoding accuracy on novel combinations of motion direction and speed (Fig. 2f), far outperforming a C3D (*25*) baseline, which exhibits *>* 40% error at high velocities (Fig. 2f); 2) The **Salience RNN** generalizes robustly across variations in centroid trajectory and spatial spread *σ*, again exceeding C3D performance (Fig. 2g); 3) The **Velocity RNN** maintains high accuracy under increased input frequency and unseen speed ranges, demonstrating strong resilience to distribution shifts (Fig. 2h). These results confirm that each pretrained RNN encodes motion attributes within smooth, low-dimensional, input-driven manifolds–rings, tori, and cylinders–that are both dynamically attractive and geometrically stable under perturbations (*14*). This structure serves as a powerful inductive bias, supporting flexible interpolation of unseen inputs while avoiding the extrapolation errors common to conventional feedforward models.

### 2.4 Representation transfer to real-world tasks for domain adaptation

Crucially, the manifolds (**input-driven continuous attractors**) learned via task-agnostic pretraining can be transferred directly to real-world scenarios, preserving both their geometric structure and dynamical properties in new tasks. This supports our core claim that these attractors act as robust, reusable computational substrates for naturalistic motion recognition without retraining. We validated this by deploying the pretrained RNN modules on two real-world datasets: the DVS Gesture dataset and a customized RGB rehabilitation action dataset. For both, we projected the neural activity evoked by real inputs into the same PC spaces used during synthetic pretraining, allowing direct assessment of whether real task trajectories remain confined to the original manifolds. Our results confirm that the manifolds’ topology and flow dynamics persist under real conditions, satisfying the two critical conditions for effective transfer (Fig. 3): 1) **Geometric alignment:** The learned manifolds (ring, torus, cylinder) mirror the topological structure of relevant motion features in natural inputs–direction, spatial position, and velocity–providing an interpretable basis for representation; 2) **Dynamical confinement:** Real input streams generate smooth, low-dimensional flows that remain close to the pretrained manifold’s tangent space, ensuring stable and robust encoding.

**Figure 3:**
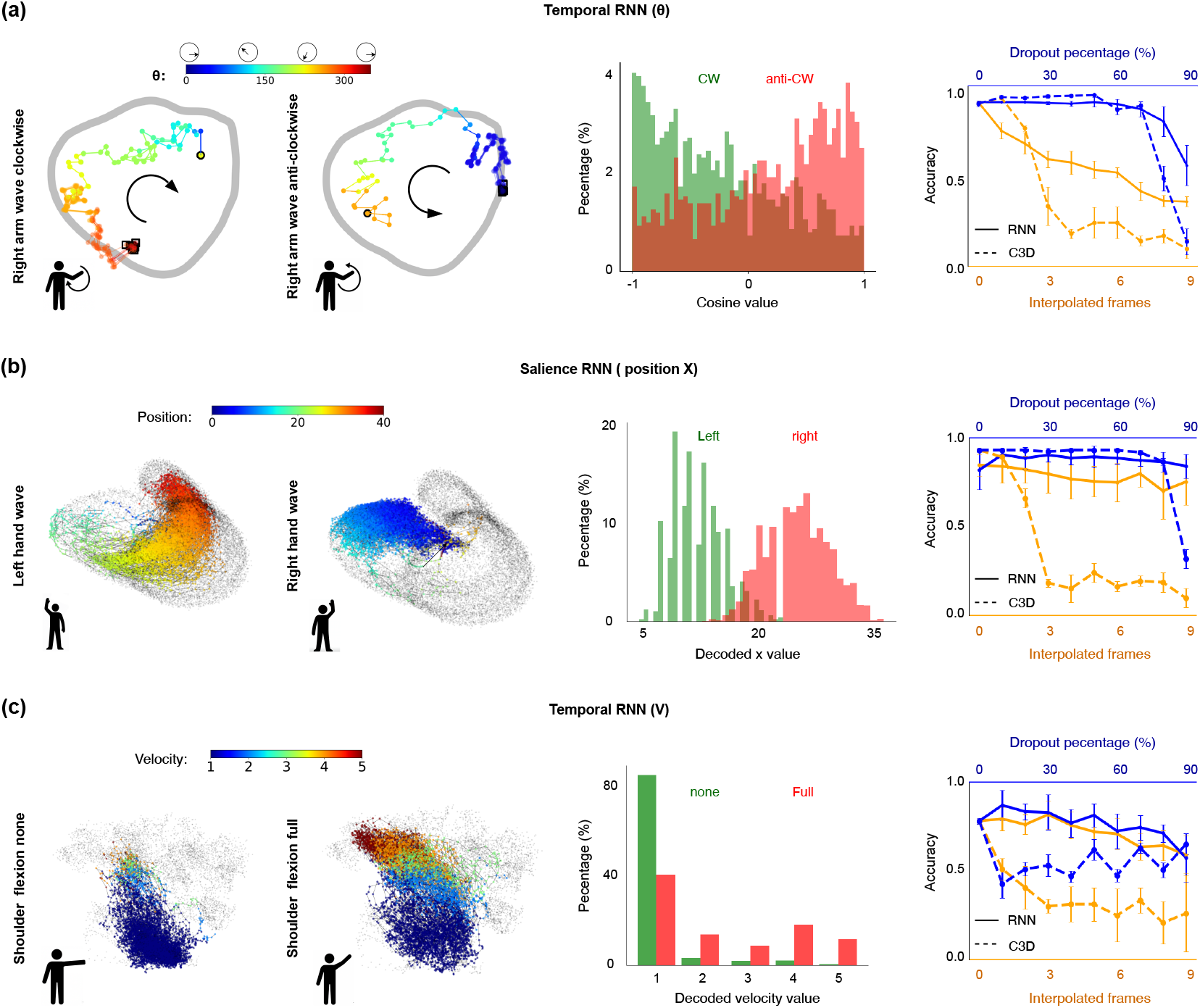
Task-specific neural dynamics and robustness of pretrained RNNs in real motion tasks. (**a**) Direction RNN (rotation *θ*): Clockwise and counterclockwise arm-waving gestures evoke distinct trajectories on the pretrained ring manifold (left); Cosine similarity between dynamics and manifold tangents confirms flow alignment (middle); Decoding remains robust under frame perturbations, outperforming C3D (right). (**b**) Salience RNN (position *x*): Left vs. right hand waves produce spatially separated trajectories on the toroidal manifold (left); Decoded *x*-positions show clear separation (middle); Decoding accuracy remains stable under input corruption (right). (**c**) Velocity RNN (speed 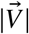): Shoulder movements of varying intensity are mapped to different regions on the velocity-tuned cylindrical manifold (left); Decoded speed levels differentiate motion intensities (middle); Decoding accuracy remains robust to temporal noise, under frame dropout/interpolation, and outperforms C3D (right).

All three RNNs transfer effectively, leveraging geometric invariance and dynamical alignment to support real-world recognition (Fig. 3): 1) **Direction RNN**: Clockwise (CW) and counterclockwise (CCW) arm-waving gestures produce distinct, oppositely circulating trajectories around the ring manifold (Fig. 3a, left). Cosine similarity confirms that neural trajectories align with the manifold’s tangent vectors, preserving directional encoding (Fig. 3a, middle). This yields near-perfect classification performance (97% accuracy) and resilience to frame dropout, outperforming the feedforward C3D (*25*) baseline (Fig. 3a, right). 2) **Salience RNN**: Left- and right-hand waves map to spatially distinct regions on the toroidal manifold (Fig. 3b, left), and decoded *x*-positions show clear separation (Fig. 3b, middle), confirming that the spatial structure learned during pretraining is preserved under naturalistic inputs. Robustness tests confirm stable decoding under noisy inputs (Fig. 3b, right). 3) **Velocity RNN**: Shoulder movements of different intensities produce distinct trajectories along the pretrained velocity-tuned cylindrical manifold (Fig. 3c, left). Decoded speed levels reliably capture motion strength, including null and full movements in the RGB rehabilitation actions (Fig. 3c, middle), and remain robust to temporal perturbations (Fig. 3c, right).

These results demonstrate that the attractors’ low-dimensional structure and input-driven dynamics enable real-time tracking, noise tolerance, and robust few-shot transfer–even when the training data is purely synthetic. Compared to static feedforward models like C3D, our RNNs continuously integrate information over time, recover rapidly from missing or corrupted frames, and preserve coherent encoding under distribution shifts (fig. S2). Our results show that **input-driven continuous attractors** provide a concrete, biologically inspired mechanism for generalizable representation transfer. They maintain topological invariance, enforce smooth flow alignment, and enable robust performance in nonstationary, noisy real-world conditions–highlighting their potential as a foundation for resilient embodied AI.

### 2.5 Brain-inspired Modular System for Interpretable Gesture Recognition

#### 2.5.1 Our PRG framework

Inspired by core principles of cortical organization, we propose a modular architecture (Fig. 4a) that integrates the motion-related attributes effectively captured by our pretrained RNN modules to enable robust, interpretable, and adaptive gesture cognition across tasks. The system begins with an input alignment module, which merges data from diverse sources into a unified binary video format at multiple spatial resolutions (**st1, st2**) (Fig. 4b), ensuring compatibility with the pretrained modules. These aligned streams are then processed by specialized feature encoding and decoding components. The overall design of this modular system draws on several well-established neurobiological principles:

**Figure 4:**
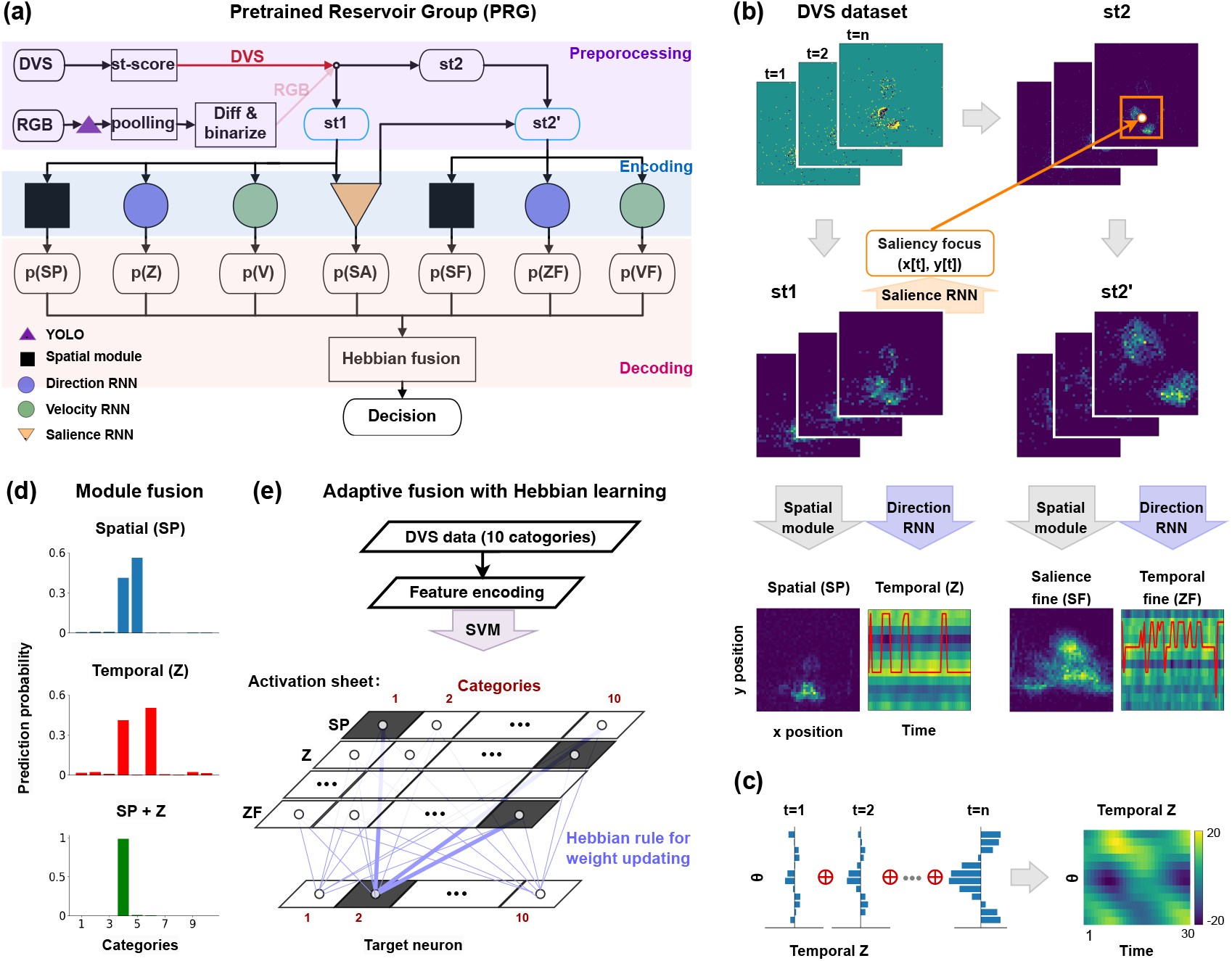
Adaptive fusion and decoding with Hebbian learning. (**a**) Overview of the PRG frame-work. (**b**) DVS input processing. At each time point *t* = 1, …, *n*, preprocessed DVS frames (**st1**) drive the salience RNN to compute the focus points (*x* [*t*], *y* [*t*]), used to crop a focused event frame (**st2**^′^). Both **st1** and **st2**^′^ are passed to downstream RNN modules to extract spatial, temporal, direction and velocity features. (**c**) Temporal encoding in the Z module. Motion directions *θ* are binned into histograms over time and concatenated into a time-by-direction map to capture the direction evolution. (**d**) Module predictions: Each RNN module outputs a probability distribution, which is combined via element-wise multiplication to sharpen and improve the final category prediction. (**e**) Hebbian-based adaptive fusion: Module outputs form an “activation sheet” (rows = modules, columns = categories). Hebbian updates (blue arrows) strengthen connections between active modules and the correct output, improving fusion over time.

- **Parallel Feature Encoding by Distinct Cortical “Mini-Columns”:** In the primary visual cortex (V1), neurons are organized into mini-columns, each selectively tuned to specific low-level visual features (e.g., orientation, spatial frequency, or motion direction (*26, 27*)). In extra-striate areas (e.g., middle temporal area (MT)/V5), neurons are further specialized for motion velocity (*28*). Mirroring this parallel architecture, our framework uses multiple RNN modules that process shared input streams, each tuned to distinct motion features: 1) **Spatial Module**: A recurrent unit with strong self-connections (autapse-like), which accumulates pixel-wise activity over time to represent spatial occupancy patterns. 2) **Direction RNN**: Classifies motion direction into 12 angular bins over a temporal window, producing coarse directional features (*Z*_+_, *Z*_−_) (Fig. 4c). 3) **Velocity RNN**: Classifies motion speed into 5 discrete speed levels, capturing trajectory-based kinematic information (*V*_+_, *V*_−_).
- **Attentional Gating Inspired by the Frontal Eye Field (FEF):** The FEF directs attention toward behaviorally relevant visual stimuli, enhancing signal fidelity while filtering out noise (*29, 30*). Analogously, our **Salience RNN** predicts the motion centroid (*x* (*t*), *y* (*t*)), which is then used to crop a high-resolution patch **st2**^′^ for fine-scale processing (Fig. 4b). This focused patch is re-analyzed by the **Direction** and **Velocity RNNs**, yielding refined direction (*Z*_+f_, *Z*_−f_) and velocity (*V*_+f_, *V*_−f_) estimates (Fig. 4b).
- **Evidence Accumulation and Bayesian-Inspired Fusion (Parietal and Prefrontal Cortex Analogy):** In biological decision-making, heterogeneous sensory cues are integrated over time via probabilistic accumulation in parietal and prefrontal circuits (*31, 32*). We implement a two-stage fusion strategy mimicking this architecture: **1) Static Fusion** (Fig. 4d): Each module’s hidden state is mapped to a class probabilities via a linear SVM. These outputs are aggregated uniformly to simulate unweighted evidence pooling. **2) Adaptive Fusion via Hebbian Learning** (Fig. 4e): A Hebbian-inspired gating layer dynamically reweighs module outputs using a small validation set, reinforcing associations between activated modules and the correct label. This allows for rapid few-shot adaptation without full retraining.

#### 2.5.2 Performance and ablation studies across two datasets

We evaluated the full PRG system on two datasets: the DVS gesture dataset and a custom RGB-based rehabilitation action dataset. The rehabilitation dataset comprises 15 fine-grained action classes involving upper and lower limb joints (e.g., shoulder, elbow, and knee), designed to simulate realistic rehabilitation scenarios with diverse camera angles and subject variability. The results demonstrate that our modular, biologically inspired model with ∼ 5-million (5M) parameters delivers high accuracy, interpretability, and robustness, even in data-limited settings. Remarkably, this compact architecture matches or outperforms much larger baselines such as a 10- or 50-layer ResNet C3D (*25*) and ViViT (*33*) models, while requiring only minutes of training time (versus hours) and using an order of magnitude fewer parameters (5M vs. 240 ∼ 300M).

On the DVS dataset (Tab. 1; Fig. 5a), our model achieves 94% accuracy across 10 gesture classes. The ablation studies illustrate each module’s role: 1) The **Spatial Module** alone achieves 80% accuracy by capturing location cues (e.g., left vs. right hand waves); 2) Adding the **Direction RNN** (Temporal *Z*) raises accuracy to 88% by encoding coarse trajectory directionality (e.g., CW vs. CCW rotations in classes 4&5); 3) Incorporating the **Salience RNN** and fine-grained features further improves accuracy to 94%, enabling the system to resolve subtle gesture variations (e.g., amplitude differences in classes 1&9). On the RGB rehabilitation action dataset (Tab. 2; Fig. 5b), our full model achieves 90% macro-class accuracy and 62% overall subclass accuracy across all 15 categories, substantially outperforming both the ResNet C3D (10- and 50-layer versions) and ViViT baselines. The ablation results show: 1) The **Spatial Module** alone achieves 43% overall accuracy with 79% macro-class accuracy and 55% averaged subclass accuracy; 2) Adding the **Velocity RNN** (Temporal *V*) improves performance to 56.5% overall with 85% macro-class and 66% subclass accuracy; 3) Incorporating the **Salience RNN** and fine-detail refinements raises performance to 62% overall with 90% macro-class and 71% subclass accuracy. Remaining fine-grained misclassifications (e.g., elbow vs. knee motions) largely stem from the inherent difficulty of projecting 3D joint dynamics onto 2D video under diverse viewpoints and distances. Nonetheless, the system robustly distinguishes coarse categories (e.g., shoulder vs. elbow motions), making it practical for physiotherapy monitoring and other real-world applications.

**Table 1:** Performance on the DVS gesture dataset: all 10 classes, classes 4&5 and 1&9.

| Network | All (10) | 4 vs. 5 | 1 vs. 9 |
| --- | --- | --- | --- |
| End-to-End trained RNN | 0.46 | 0.50 | 0.55 |
| C3D(10) | **0.94** | **0.98** | 0.81 |
| ViViT | 0.93 | 0.87 | 0.51 |
| Temporal Z | 0.66 | 0.90 | 0.70 |
| Temporal Z + Spatial | 0.88 | 0.95 | 0.83 |
| Our model: Temporal Z+ Spatial + Saliency | **0.94** | **0.98** | **0.85** |

**Table 2:** Performance on the rehabilitation RGB dataset.

| Action type | OUR MODEL | C3D(10) | C3D(50) | ViViT |
| --- | --- | --- | --- | --- |
| Macro class | **0.90** | **0.90** | 0.86 | 0.75 |
| action 1 | **0.81** | 0.58 | 0.56 | 0.44 |
| action 2 | **0.81** | 0.55 | 0.49 | 0.50 |
| action 3 | **0.78** | 0.40 | 0.30 | 0.41 |
| action 4 | **0.63** | 0.32 | 0.32 | 0.35 |
| action 5 | **0.49** | 0.44 | 0.41 | 0.43 |
| overall (15 classes) | **0.62** | 0.46 | 0.43 | 0.36 |

**Figure 5:**
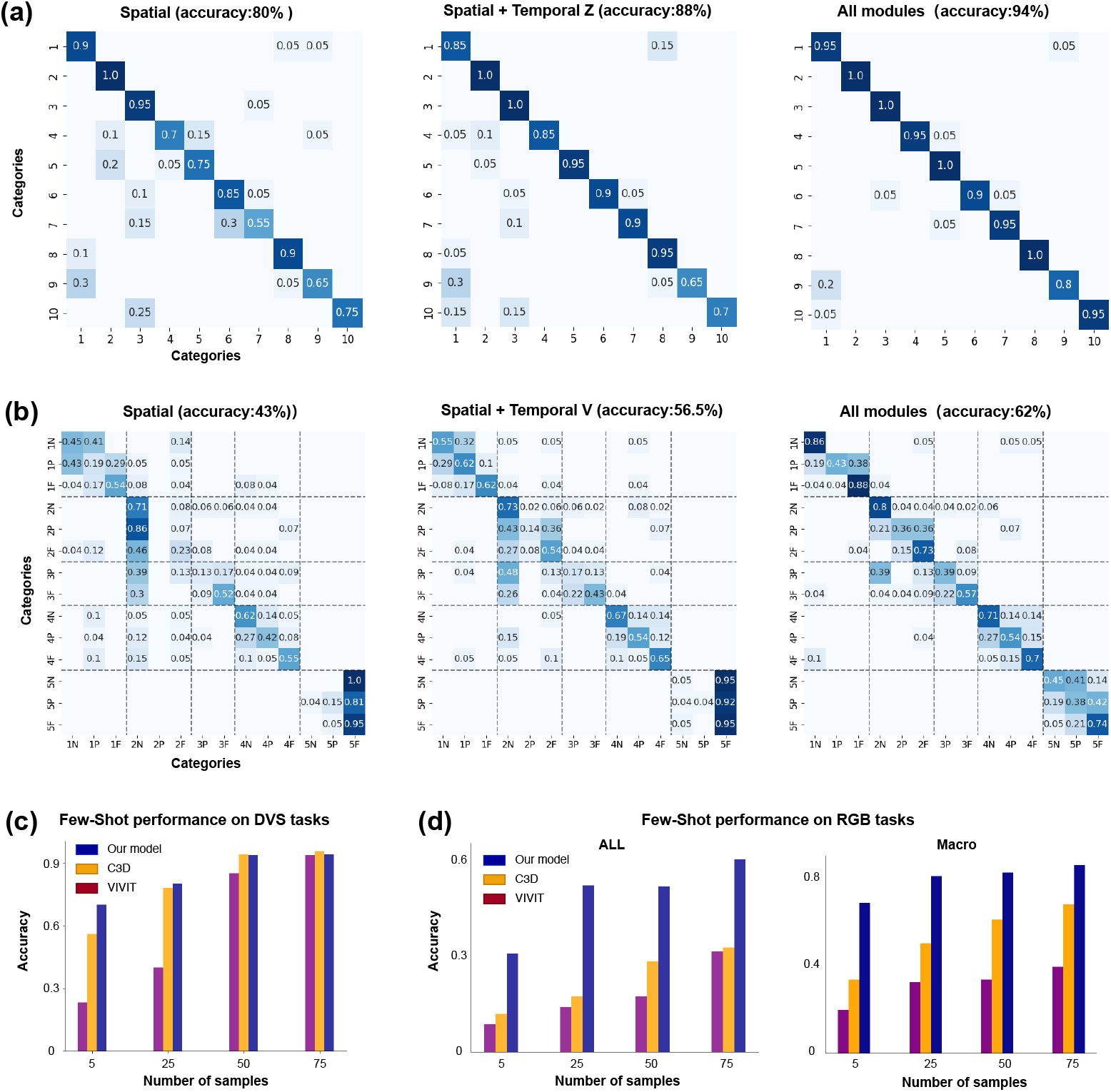
Model performance on DVS and RGB tasks. (**a**) DVS task–Confusion matrices: Comparison of classification accuracy using spatial module only (left), spatial + temporal(velocity+direction) modules (middle), and all modules (right). Darker colors indicate better performance. (**b**) RGB task– Confusion matrices: Comparison of classification accuracy using spatial module only (left), spatial + temporal modules (middle), and all modules (right). Darker colors indicate better performance. DVS task–Few-shot learning: Our model (blue) outperforms C3D (orange) and ViViT (purple) in low-data regimes. (**d**) RGB task–Few-shot learning: Bar plots of overall (left) and macro-averaged (right accuracy for different sample size. Our model (blue) consistently outperforms C3D (orange) and ViViT (purple), especially in low-data regimes.

In few-shot scenarios, our system demonstrates strong sample efficiency and adaptability: 1) **DVS dataset** (Fig. 5c): With only 5 samples per class, the full model achieves ≈ 70% accuracy, substantially outperforming 3D ResNet-50 (58%) and ViViT (28%); 2) **RGB dataset** (Fig. 5d): At just 5 samples per class, our model attains ≈ 75% macro-class accuracy, whereas the ResNet C3D (10 layers) and ViViT achieve only ≈ 20% and ≈ 21%, respectively. The macro-class accuracy steadily increases, reaching nearly 80% as the number of shots grows, with overall accuracy following a similar trend. These results highlight the benefits of modular pretraining and Hebbian-style adaptive fusion for low-resource gesture recognition, demonstrating strong few-shot learning capability and continual adaptability in real-world rehabilitation and monitoring contexts.

#### 2.5.3 Interpretability of Modular Representations

To verify that the transferred representations remain interpretable in downstream tasks, we conduct fully unsupervised kernel and hierarchical clustering analyses (see Sec. 5 for details) to examine how individual modules encode meaningful features across tasks. These analyses reveal that each module maintains distinctive, reusable representational structures from pretraining through deployment. In the DVS task (Fig. 6a), the **Spatial module** partitions the scene into task-relevant regions (e.g., left/center/right or upper/lower fields), while the **Spatial fine module** further resolves localized and subtle variations–such as distinguishing left-hand from right-hand waves–resulting in compact, well-separated clusters. Meanwhile, the **Direction module** (temporal *Z*) captures global motion patterns, such as CW vs. CCW rotations. These distinctions are evident in the clear cluster boundaries in the dendrograms (Fig. 6a) and the consistent separations in the 2D embeddings (Fig. 6b). In the rehabilitation task (Fig. 6c-f), the **Salience, Spatial** and **Velocity** (temporal *V*) modules each separate action 1 apart from actions 3, 4, 5 (Fig. 6c), by leveraging different features–such as knee movements captured by the **Spatial module** (Fig. 6e). Additionally, the three temporal modules (*V, Z*, and *V F*) characterize subclasses within action 1 (Fig. 6d), cleanly distinguishing low (None) vs. high (Full) movement intensities in the 2D embeddings (Fig. 6f). This clear, modular “alphabet” of motion primitives–absent in monolithic black-box networks–underscores the interpretability and transferability advantage of our framework.

**Figure 6:**
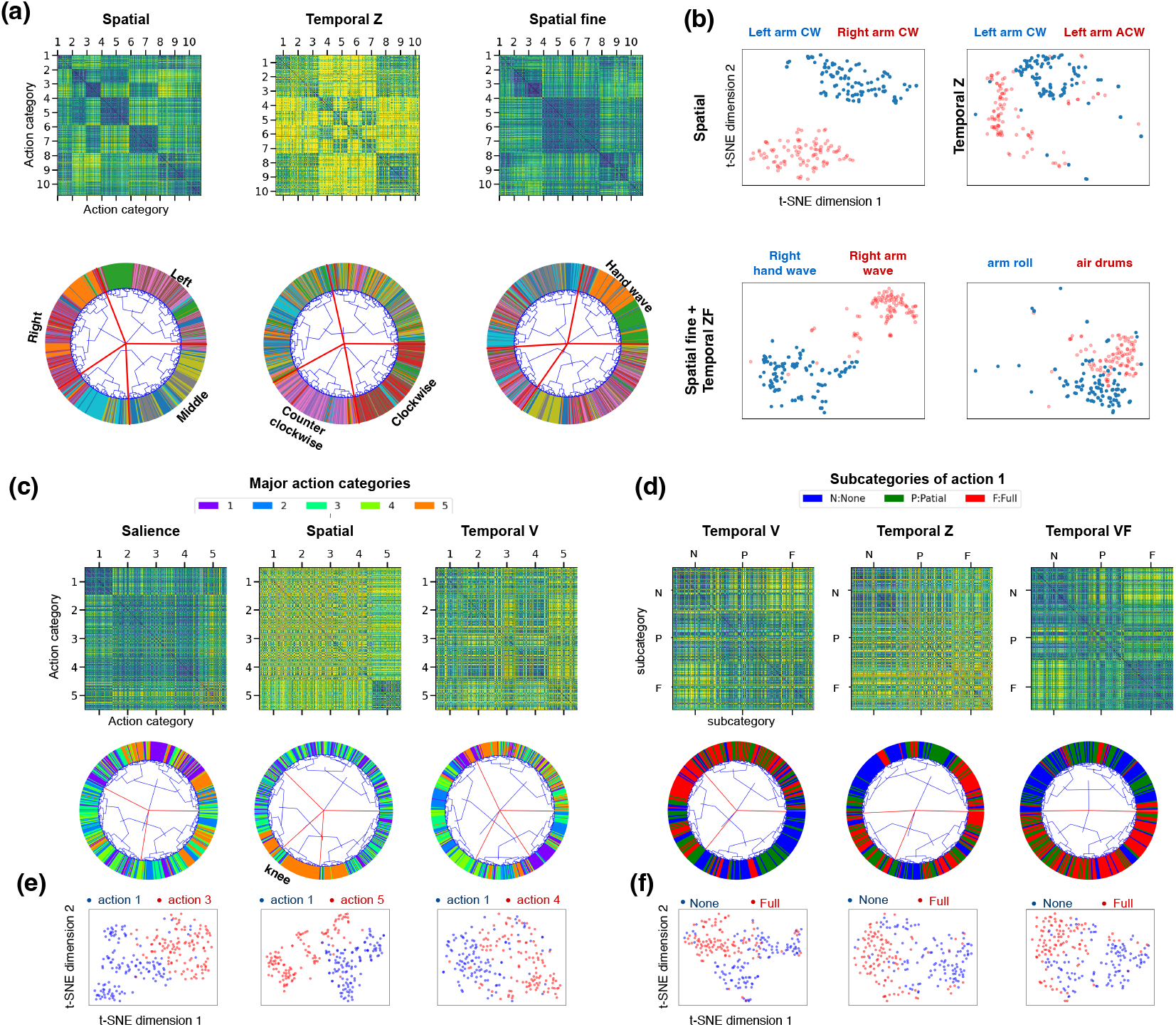
Module-specific Representations across tasks (DVS task: (**a**-**b**); RGB task: (**c**-**f**)). (**a**) Kernel matrices (top) show pairwise distances between internal states for each module (spatial (SP), temporal (T), spatial fine (SF)) across 10 action categories, revealing distinct representational patterns; Circular dendrograms (bottom) display how each module clusters actions, capturing structure such as left/right arm movements, motion direction (CW/CCW), and hand-related actions. Cluster labels summarize shared features within groups. (**b**) t-SNE plots show clear separations among actions like “Left arm CW”, “Right arm CW”, and “Left arm CCW”, reflecting strong spatial and temporal selectivity. (**c**) Kernel and clustering analysis across 5 major action categories. (**d**) Kernel and clustering analysis across 3 fine-grained subcategories. (**e**) t-SNE plots showing clear separations among different macro action groups for each module. (**f**) t-SNE plots showing clear separations among different micro action groups for each module.

## 3 Discussions

We have presented a biologically inspired modular RNN architecture where a task-agnostic pretraining sculpts reusable **input-driven continuous attractor manifolds**, which confer strong resilience to substantial input perturbations (e.g., up to 50% frame dropout). These pretrained, invariant input-driven manifolds can be effectively transferred to real-world tasks, enabling our framework to match or surpass the performance of conventional end-to-end deep networks while using far fewer parameters across diverse tasks and sensor modalities (e.g., DVS and RGB). Each RNN module encodes a specific low-dimensional manifold aligned with key sensorimotor primitives–direction, velocity, or position–mirroring computations found in cortical circuits; Our findings suggest that many sensorimotor recognition tasks can be reframed as trajectories on shared, reusable low-dimensional manifolds, offering an efficient and robust alternative to monolithic deep networks. Our contributions lay the groundwork for self-configuring, resilient agents capable of real-time learning and adaptation with minimal data and computational overhead.

### 3.1 Representation Transfer for domain adaptation

Domain adaptation (*2,34,35*) remains a defining challenge for modern AI, yet it is a natural strength of biological systems. Machine learning systems typically extract narrow, task-specific regularities, whereas robust generalization requires features that remain invariant for effective transfer across domains. To approximate such invariance, prevailing AI methods depend heavily on large-scale data augmentation (*36*), contractive autoencoders (*37*), adversarial training (*38–40*), or causal representation learning (*41*). While these techniques can be powerful, they are often compute-intensive, data-hungry, and yield opaque, black-box features that are hard to interpret, control, or reuse. Reservoir computing (*4, 42*) offers a contrasting idea: instead of end-to-end training for every task, it uses a fixed, randomly connected RNN that projects temporal inputs into a high-dimensional dynamical reservoir, training only a simple readout layer for task-specific output. This approach dramatically reduces training cost and data requirements for time-series tasks. However, as the reservoir’s internal dynamics are random rather than structured with feature learning, its capacity to handle complex, highly structured sensory streams like videos is limited–it lacks inductive biases needed for more sophisticated, reusable representations.

Biological brains, by contrast, achieve efficient domain adaptation without large labeled datasets, partly because they reuse highly structured, evolutionarily conserved circuits across tasks (*43, 44*). Unlike reservoir networks, these neural circuits are not random (*45,46*), but evolutionarily organized into modular architectures that encode stable, reusable priors–what cognitive science refers to as basic concepts (*47,48*). For example, grid cells provide a flexible spatial metric that mammals reuse for non-spatial cognitive reasoning beyond navigation (*49, 50*).

Our framework directly leverages this biological principle. Instead of relying on purely task-specific representations, we anchor RNN modules in kinematic primitives—direction, velocity, and position—that are fundamental to motion perception in the brain (e.g., MT/V5 neurons) (*19, 28*). These variables act as robust, interpretable building blocks that naturally remain invariant across a wide range of tasks and conditions. Here we used a simple curriculum learning protocol to embed these invariances into neural dynamics, by pretraining each module on synthetic proxy tasks. These tasks isolate individual motion attributes using carefully controlled symmetries and uniform sampling, shaping each RNN into a compact, input-driven continuous attractor. This pretraining sculpts low-dimensional manifolds that are smooth, interpretable, and aligned with fundamental physical symmetries–structures that real-world data alone rarely expose cleanly. Our approach eliminates the need for extensive labeled data, expensive augmentation pipelines, or heavy adversarial schemes (*36, 38*), by focusing on synthetic, well-defined motion sequences instead of massive real-world datasets.

The resulting modules remain lightweight, transparent, and easy to compose: they can be rapidly adapted for new tasks with minimal additional training while preserving clear, physically grounded interpretability. By grounding representation transfer in evolutionarily conserved kinematic primitives and embedding them as reusable input-driven attractors, our framework offers an interpretable, data-efficient alternative to black-box domain adaptation methods. This biologically inspired approach closes the gap between generalization power and practical deployment, pointing toward more robust, energy-efficient AI that mirrors how brains adapt in an unpredictable world.

### 3.2 Invariant Input-driven Neural Manifolds

A central insight of our work is the discovery of **input-driven continuous attractors** in pretrained RNNs–an important mechanism for robust representation transfer and reliable neural information processing via high-dimensional, input-induced transient dynamics (*51*). Unlike classic autonomous attractors (*52*), which settle into fixed states and stay there once input is removed, our attractors remain continuously driven by external sensory streams. When input features are constant, the system settles into a stationary region with minimum flow, behaving like a classical point attractor; when the input varies, however, it generates smooth, low-dimensional flows that track the changing input in real time. In doing so, it transforms noisy, high-dimensional signals into robust, interpretable trajectories on low-dimensional manifolds embedded within a high-dimensional neural state space. This dynamical behavior extends classic continuous attractor neural networks (*53*), which typically implement low-dimensional autonomous dynamics for tasks like spatial integration. Our input-driven attractors uniquely combine three critical dynamical properties: 1) **Stability**: robust fixed-point convergence when input is static; 2) **Adaptability**: continuous tracking of varying input; 3) **Flexibility**: compact, low-dimensional manifolds embedded within a high-dimensional hidden state space. This unique combination distinguishes our findings and enables the learned manifolds to transfer effectively from simple synthetic pretraining tasks to complex, real-world conditions, supporting strong OOD generalization.

While structured manifolds can, in principle, also appear in purely feedforward architectures (e.g., sparse autoencoder (*54*), CNNs (*55, 56*)) under carefully designed training protocols, these models perform static input-output mappings and lack our three unique dynamical properties. As a result, feedforward networks processing sequential input (e.g., videos) must rely on fixed-length windows, which limits temporal continuity and makes them brittle to perturbations and frame dropout, as demonstrated in Fig. 3. By contrast, our input-driven continuous attractors maintain coherent low-dimensional internal dynamics that adapt in real time, keeping representations stable and interpretable even under severe disruptions (e.g., high frame dropout or domain shifts) and enabling few-shot adaptation with minimal extra data or computation.

Strikingly, the geometries we observe in these invariant input-driven continuous attractors echo patterns in biological circuits, such as the toroidal manifolds formed by grid cells in the rodent entorhinal cortex (*7*) or ongoing movement signals during decision making of anterior cingulate cortex neurons (*57*). This parallel suggests that embedding simple, low-dimensional manifolds within high-dimensional recurrent dynamics may be a fundamental neural strategy for balancing stability with real-time flexibility–an insight that underpins our framework’s robust representation transferability and adaptability in dynamic real-world conditions.

### 3.3 Advantages of Brain-inspired Framework

Human-like transfer learning relies heavily on the principle of recomposition–combining existing functional building blocks to tackle new tasks (*58, 59*). Our framework extends this principle by designing a modular architecture inspired by cortical microcircuits: parallel processing streams mimic minicolumns, while a salience mechanism supports multi-resolution perception and flexible integration of diverse input sources. This allows the system to decompose incoming sensory data into interpretable “symbols” via pretrained, invariant input-driven neural manifolds, then flexibly recombine these primitives for new tasks.

A practical advantage is that when facing a novel task, only a lightweight decoder needs to be updated–dramatically reducing training time, data transfer, and computational cost. This plug-and-play modularity makes robust inference feasible on low-power hardware such as FPGAs or neuromorphic chips, while keeping raw sensory data local to preserve privacy and reduce latency. As a result, our architecture naturally supports the demands of edge computing and embodied intelligence, paving the way for self-configuring agents that can learn and adapt in a continuous manner to the unpredictable physical world.

While our demonstration focused on relatively simple rigid-body-like actions, the same principles can be extended to more complex behaviors by adding new modules to the existing framework. As these extensions reuse the same recomposable attractor primitives, the cost of expanding capabilities remains low. For example, a segmentation module could first partition complex scenes into distinct regions, while chain-attractors or sequence-memory motifs could encode multi-step or hierarchical tasks. Each extension preserves interpretability and transferability by reusing the same pretrained manifolds, minimizing costly retraining while expanding the agent’s behavioral repertoire.

## 4 Summary

In summary, our framework combines the stability of classical attractor networks with the adaptability of input-driven RNNs through task-agnostic pretraining. We sculpt invariant input-driven continuous attractors that maintain stable, low-dimensional representations even under abrupt input changes, enabling robust representation transfer and few-shot adaptation in nonstationary, real-world settings. This biologically inspired modular design also allows agents to expand their attractor repertoire through synthetic pretraining–using, for example, generative models or physics-based simulators–and to dynamically select and compose modules via meta-learning or neural architecture search. Together, these capabilities lay the groundwork for resilient, low-cost embodied AI that can learn continually and adapt in real time with minimal data and computation.

## 5 Methods

### 5.1 RNN pretraining

All three RNN modules are obtained through a task-agnostic pretraining method with synthesized input videos designed as follows. To ensure versatility across various scenarios, we employ the original vanilla recurrent neural network (RNN) with 512 neurons, with an input weight matrix *W*_*I*_, an internal matrix *W*_rec_, and a readout matrix *W*_*O*_. The decay timescale *τ* is set to be 2 time steps. The internal matrix *W*_rec_ is initially a sparse matrix with 80% positive and 20% negative values, mimicking the E-I network in the cortex (*60*).

The hidden state *h*(*t*) in the RNN evolves as:

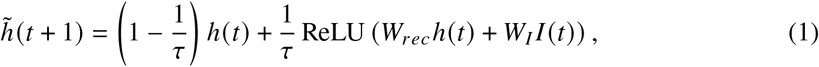

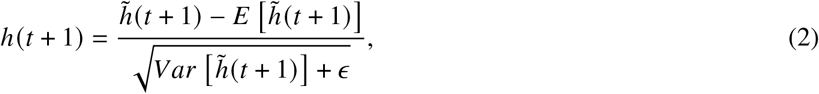

where *I* (*t*) represents the input frame at time *t* flattened into a 1D vector, *ϵ* is a small positive offset, and *E* [·] (*V ar* [·]) denotes the mean (variance) of the signal. To maintain consistency across different input conditions and enhance network stability, layer normalization (Eq. 2) is applied at each time step, ensuring that the hidden state *h*(*t*) remains normalized throughout the network’s operation.

A linear readout function and cross-entropy loss are used to train the network:

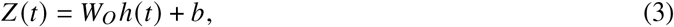

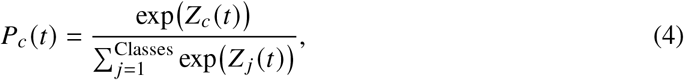

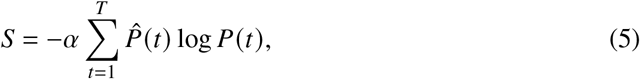

where 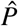 is the one-hot ground truth label, *P* is the estimated probability, Classes is the number of categories, *T* is the total time steps per clips. The optimization uses truncated backpropagation through time (*61*) with an Adam optimizer at a learning rate of 10^−4^. The input video and training target vary across the direction, salience, and velocity networks, and their respective network configurations are provided in Tab. 3. All models converge within 10^6^ training episodes, requiring approximately 48 hours of computation on a single NVIDIA A40 GPU.

**Table 3:** Pretraining configurations and readout functions for recurrent neural networks.

| Network | Category | Parameter | Value |
| --- | --- | --- | --- |
| Direction RNN | Pretraining | Velocity range | 0–1(pixel per frame) |
|  |  | Clip length | 100 |
| | | Noise model | Bernoulli($p = 0.01$) |
| | | Prediction target | Motion direction $\theta$ |
| | Readout | Output variable(s) | $Z(t)$ |
|  |  | Classes | 12 |
| | | Loss function | $S(\text{cross entropy})$ |
| Saliency RNN | Pretraining | Velocity range | 0–1 |
|  |  | Clip length | 50 |
| | | Noise model | Gaussian($\sigma \in [2, 8]$) |
| | | Prediction target | Centroid $(x, y)$ |
| | Readout | Output variable(s) | $Z_x(t), Z_y(t)$ |
| | | Classes | $42(x), 42(y)$ |
| | | Loss function | $S_x + S_y$ |
| Velocity RNN | Pretraining | Velocity range | 1–5 |
|  |  | Clip length | 50 |
| | | Noise model | Bernoulli($p = 0.01$) |
|  |  | Prediction target | Speed, direction |
| | Readout | Output variable(s) | $Z_v(t), Z_\theta(t)$ |
| | | Classes | $5(v), 12(\theta)$ |
| | | Loss function | $S_v + S_\theta$ |

### 5.2 Data Acquisition and Preprocessing

We evaluate our system using two distinct datasets. The DVS Gesture Dataset (*62*) provides event-based recordings particularly suited for motion recognition tasks. Additionally, we curate a custom RGB-based rehabilitation action dataset inspired by the Fugl-Meyer Assessment (*63*), consisting of 1,678 samples divided into five major categories (Fig. S1):

1. Shoulder flexion from 90^°^ to 180^°^
2. Shoulder flexion from 0^°^ to 90^°^
3. Shoulder abduction from 0^°^ to 90^°^
4. Pronation and supination
5. Knee flexion to 90^°^

Each category is further subdivided into three subcategories based on recovery levels: none (0), partial (1), and full (2). The gestures are recorded using an RGB camera from various viewing angles and distances to simulate real-world usage scenarios. A summary of the two datasets is provided in Tab. 4.

**Table 4:** Quantitative summary of datasets.

| Dataset | Total samples | Total classes | Samples per class | Train/Test split |
| --- | --- | --- | --- | --- |
| DVS | 1,220 | 10 | 122 | 4:1 |
| RGB | 1,678 | 15 (5 macro×3 levels) | 112 | 4:1 |

To ensure compatibility with both DVS and RGB streams, we design a unified preprocessing pipeline that (1) converts inputs into a standardized binary video format and (2) denoises in both spatial and temporal domains. For DVS data, we employ a dedicated spatio-temporal core (ST-core) structure (*64*) for noise reduction (Fig. 4(b)). The ST-core consists of binary neural networks that perform spatial and temporal processing, ensuring effective denoising while preserving critical spatio-temporal information. The spatial computation integrates the data as follows:

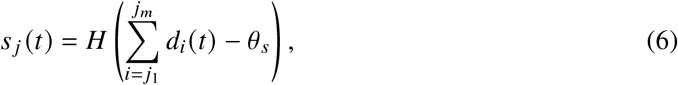

where *d*_*i*_ (*t*) is the value of the *i*^th^ pixel at time *t, s* _*j*_ (*t*) is the output pixel, and integration occurs over a detection range of square region *m* = Δ*ST*_*s*_^2^(correspond to *j*_1_ to *j*_*m*_ pixel in input frame). The Heaviside step function *H* (*S* − *θ*_*s*_) equals 1 if the sum *S* exceeds the spatial threshold *θ*_*s*_, and 0 otherwise. Temporal computation is performed as:

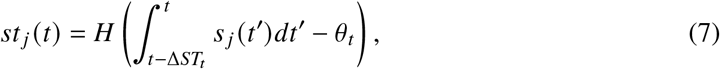

where Δ*ST*_*t*_ is the temporal integration window and *θ*_*t*_ is the temporal threshold. The parameters Δ*ST*_*s*_, *θ*_*s*_, Δ*ST*_*t*_, *θ*_*t*_ control the spatial and temporal receptive fields and thresholds of the ST-core. We generate two binary video resolutions: **st1** (42 × 42) with parameters 3, 2, 3, 2 and **st2** (128 × 128) with parameters 1, 1, 2, 2.

For RGB video preprocessing, we first employ YOLOv8 (*65*) to detect the subject and extract a 500 × 500 region centered on the detected bounding box from the original 1920 × 1080 video. To generate multiscale inputs consistent with the DVS modality, we apply average pooling with window sizes of 10 × 10 and 4 × 4, producing two spatial resolutions referred to as **st1** and **st2**.

To capture motion dynamics and align with the sparsity characteristics of DVS input, we compute the pixel-wise temporal difference between each frame and the final frame of the clip. The resulting difference maps are then binarized using a fixed threshold (95%), yielding binary video representations that mimic the temporal sparsity of event-based data.

### 5.3 Feature Encoding

Our modular, pretrained RNNs extract 11 complementary feature streams from each input sequence, encompassing spatial layout, motion dynamics, and salience trajectories (Fig. 4, S1).

#### Spatial Distribution

We first collapse temporal dynamics to capture the overall spatial layout.

For each pixel *j*, we sum the low-resolution binary frames **st1**(*t*) over time:

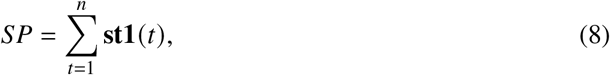

yielding a 2D activity map that encodes zero-order spatial features, reflecting the cumulative spatial distribution of events (Fig. S1).

#### Bidirectional Motion Code

To capture bidirectional motion patterns, we feed both the original and time-reversed low-resolution sequences into pretrained Direction and Velocity RNNs, yielding readout vectors for direction [*z*(*t*), *z*_−_(*t*)] and velocity [*v* (*t*), *v*_−_(*t*)]. Concatenating across frames produces global motion codes S1:

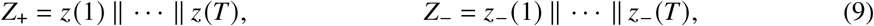

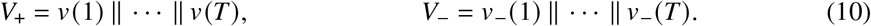

This bidirectional temporal code summarizes the evolution of motion direction and speed across time, providing a compact representation of global motion dynamics.

#### Salience Trajectory

The Salience RNN tracks the spatiotemporal focus of motion over time, outputting probability distributions *z*_*x*_ (*t*) and *z*_*y*_ (*t*) over horizontal and vertical positions, respectively. These are concatenated into per-frame vectors *f* (*t*) = *z*_*x*_ (*t*), ∥, *z*_*y*_ (*t*), which are then sequentially aggregated to form a trajectory (Fig. S1):

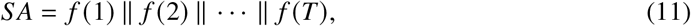

resulting in a high-dimensional trajectory that captures attentional shifts throughout the sequence. **Fine-Detail Encoding** To capture subtle local variations, we leverage high-resolution input frames **st2**. Using the predicted salience center 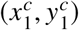 from the low-resolution stream **st1**, we scale the coordinates to the **st2** resolution as:, 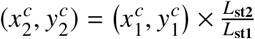, and extract a centered crop from **st2**, denoted as **st2**^′^. We then apply the same RNN-based feature extractors to **st2**^′^ to obtain fine-grained spatial features (*SF*), as well as refined direction and velocity codes (*Z F*_+_, *Z F*_−_, *V F*_+_, *V F*_−_).

These features are particularly effective for disambiguating gestures that share similar global patterns but differ in localized movements.

Collectively, the eleven feature descriptors {*SP, Z*_+_, *Z*_−_, *V*_+_, *V*_−_, *S A, SF, Z F*_+_, *Z F*_−_, *V F*_+_, *V F*_−_} constitute a compact yet expressive representation for general-purpose action encoding.

### 5.4 Feature Decoding

The decoding pipeline comprises two sequential stages: per-module probabilistic mapping and modular fusion. This design combines efficient SVM-based classification with a biologically inspired evidence integration strategy (Fig. 4).

#### 5.4.1 Stage I: Per-Module Probabilistic Mapping

For each descriptor, a Support Vector Machine (SVM) with a Radial Basis Function (RBF) kernel is trained to produce a class probability vector, *p* (*X*) = SVM_RBF_(*X*), where *X* denotes the feature descriptor. The decision function is 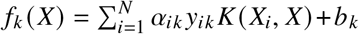, with the RBF kernel defined as 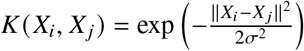.

We use Platt scaling (*66*) to convert these decision scores into calibrated probability estimates. The resulting probability that sample *i* belongs to class *k* is computed using the softmax function over the scaled decision scores:

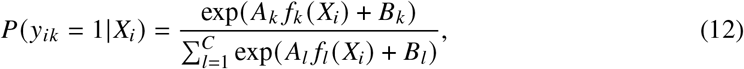

where the scaling parameters *A*_*k*_ and *B*_*k*_ are determined by maximizing the log-likelihood function on a calibration set:

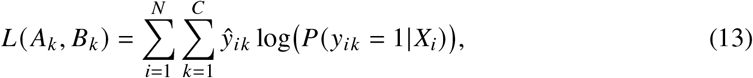

where *ŷ* _*ik*_ denotes the one-hot encoded ground-truth label for sample *i* and class *k*.

#### 5.4.2 Stage II: Modular Fusion

To obtain the final prediction, we fuse the outputs of individual modules using two alternative schemes: a simple multiplicative fusion and an adaptive Hebbian learning-based fusion.

##### Simple Fusion

Assuming equal reliability, we compute the element-wise product of all module probability vectors:

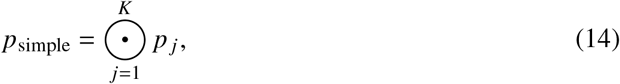

where *p* _*j*_ is the probability vector from module *j*.

##### Adaptive Hebbian Fusion

To account for variable feature importance, we first perform *module selection* by retaining only the top-*k* performing modules based on validation accuracy. We convert each module’s probability output *p* _*j*_ to a one-hot activation *B* _*j*_ via winner-take-all and feed these into a Hebbian-learning layer (*67*) with weight matrix *W* ^*H*^. The weights are updated during training:

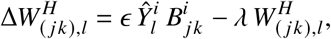

where 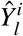 is the true class indicator, *ϵ* is the learning rate(taken as 10^−3^), and *λ* is the decay rate(taken as 10^−5^). At inference, the final weighted log-likelihood for class *l* is computed as:

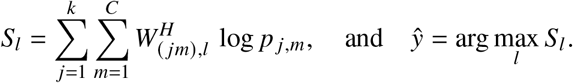

### 5.5 Slow Manifold Analysis

Recurrent neural networks (RNNs) can be viewed as nonlinear dynamical systems whose internal states tend to settle near “slow points” once trained (*14*). We empirically locate these slow-manifold regions by following a systematic process.

#### 1. Record internal states

To probe each pretrained RNN’s dynamics, we present two types of stimuli:

- *Artificial sequences:* 500 synthetically generated clips (100 frames each) covering the full range of motion parameters. This data is for manifold analysis.
- *Real sequences:* The entire training set of DVS gesture and RGB rehabilitation action videos at resolution **st**_1_. This data is to probe representation transfer.

For every frame *t*, we record the input *x*_*t*_, its ground-truth order parameter *ϕ*_*t*_, and the network’s hidden state *h*(*t*), yielding paired datasets {(*x*_*t*_, *ϕ*_*t*_, *h*(*t*))}.

#### 2. Perturbation analysis

We perform perturbation analysis on both internal states and input parameter.

- *Internal state perturbation:* For each pair (*x*_*t*_, *h*_*t*_), we apply controlled deviations, such as linear perturbation:

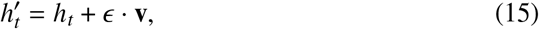

where *ϵ* ∈ ℝ is a small scalar and **v** is a unit direction vector along *h* _*t*_.
- *Input space perturbation:* We perturb order parameters such as direction, velocity, or position:

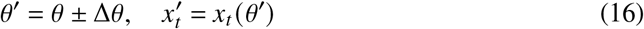

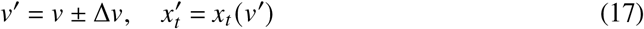

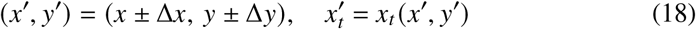

where we perturb each order parameter (e.g., direction *θ*, velocity *v*, spatial position (*x, y*)) to generate a new input 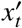 with modified stimulus properties.

#### 3. Dynamics flow record

For each state *h*_*t*_, we evaluate contractive dynamics by allowing it to evolve under a constant input *x*_*t*_ for *k* steps(taken as 10) to a relaxed state 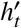. The relaxation process is:

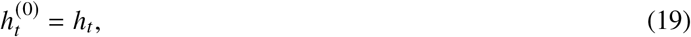

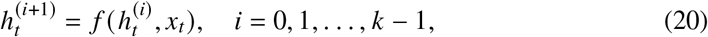

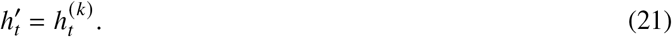

The final relaxed state 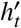 reflects the attractor tendency under constant input. We compute the average per-step change as: 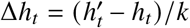 from which we extract two quantities: velocity *v*_*t*_ = ∥Δ*h*_*t*_ ∥_2_ and direction *d*_*t*_ = Δ*h*_*t*_/∥Δ*h*_*t*_ ∥_2_.

#### 4. Dimensionality reduction

We apply Principal Component Analysis (PCA) to the collected hidden states and project them into a 3D space using the first three principal components.

#### 5. Visualization

The hidden states are plotted in the 3D PCA space, colored by their corresponding order parameter. We overlay the manifold “ridge,” computed as the centroid of PC coordinates for each discrete parameter value.

#### 6. Quantifying Contraction

For each perturbed state, we compute the update vector Δ*h*_*t*_ and its direction *d*_*t*_. We then measure the angle to the manifold:

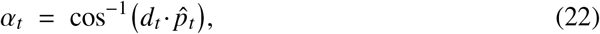

where 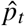 is the vector from the state to the manifold. Angles *α*_*t*_ near 180^°^ indicate a restoring pull. Plotting *v*_*t*_ versus dist(*h*_*t*_, ℳ) reveals how contraction speed varies with displacement.

### 5.6 Hierarchical Clustering Method

We analyze module representations using the hierarchical clustering method (*68*). The processed video **st1** is taken for analysis. These features are transformed by their respective modules into high-dimensional representations. To facilitate analysis, we apply t-SNE to project these representations into a low-dimensional space. We then compute pairwise distances between all data points to form a distance kernel matrix, which serves as input for bottom-up hierarchical clustering. The resulting dendrogram provides a hierarchical structure of the data, allowing us to interpret relationships between instances at different granularities.

### 5.7 C3D and ViViT Networks

To benchmark our system, we implement two representative video architectures.

#### ResNet-50-based C3D

Our 3D CNN backbone is based on an inflated ResNet-50 (*69*), where 2D convolutions are extended to 3D. Specifically, the first layer applies a 3D convolution with a kernel size of 7 × 7 × 7 and stride (1, 2, 2), followed by 3D batch normalization, ReLU activation, and a 3 × 3 × 3 max-pooling layer. All weights are initialized using Kaiming initialization. We train the model on both the DVS gesture and RGB rehabilitation datasets for 30 epochs using the Adam optimizer (initial learning rate 10^−4^, halved every 10 epochs; weight decay 10^−5^) with a batch size of 16.

#### ViViT

We implement a vanilla Video Vision Transformer (ViViT) following the “Factorised Encoder” variant proposed by Arnab et al. (*33*). This architecture is designed to efficiently capture spatio-temporal patterns in video data:

- **Input Embedding:** Each input video is divided into non-overlapping tublets using a 3D convolution with kernel and stride size of 2 × 16 × 16. Each tublet is then linearly projected into a 1024-dimensional embedding. A typical video clip is thus transformed into a sequence of 1536 tokens. Learnable positional embeddings are added to preserve spatiotemporal order before feeding the tokens into the transformer.
- **Transformer Encoder:** The encoder comprises 12 layers of factorized self-attention blocks, each consisting of pre-normalization LayerNorm, 12-head multi-head self-attention, and a feedforward MLP with GELU activation.

The model is trained for 50 epochs on both DVS and RGB rehabilitation action dataset (batch size 8) using AdamW (learning rate 5 × 10^−5^, weight decay 10^−4^).

## Acknowledgments

We thank Bailu Si and Sen Song for fruitful discussions.

## Author contributions

T.X. and D.Y. designed the study; T.X. performed the experiments and analyses; D.Y. and T.X. wrote the manuscript; All authors contributed to the discussion.

## Data and materials availability

All data needed to evaluate the conclusions in the paper are present in the paper and/or the Supplementary Materials. The custom RGB action dataset is available at https://doi.org/10.5281/zenodo.16454040 and https://doi.org/10.5281/zenodo.16473362. Computer code for all simulations and analysis of the resulting data is available at https://doi.org/10.5281/zenodo.16441066.

## Supplementary materials

Supplementary Note 1 and Figures S1 to S2

## Supplementary Materials for

### Supplementary Note 1

#### Reservoir Computing (RC)

RC leverages the inherent chaotic dynamics of large, randomly connected RNNs to process complex temporal patterns. Unlike traditional deep learning, RC does not require training of recurrent weights, significantly reducing computational complexity and data requirements. Recent studies show that such randomly initialized networks can retain high expressive power, enabling effective transformations of input data into high-dimensional representations (*70*). This makes RC particularly appealing in scenarios where deep learning implementations are constrained by resources or data scarcity (*71, 72*).

#### Spike Gating Flow (SGF)

SGF (*64*) is a few-shot learning framework based on the Neural Engineering Framework (NEF) (*73*), designed to embed prior knowledge in a hierarchical architecture with three guiding principles: 1) Emphasis on global features of movement rather than local pixels; 2) Pre-designed hierarchical modules, each specialized for particular spatiotemporal patterns; 3) Histogram-based training that adjusts output weights using global feature history. The SGF system matches or surpasses LSTM (*74*) performance with significantly fewer samples and lower energy usage. In contrast, our framework here improves task performance (from 87.5% to 94%), reduces dependency on hand-crafted components, and enhances robustness to noise, making it applicable across a wider range of real-world tasks.

#### Brain-inspired Models for Spatiotemporal Pattern Recognition

Inspired by subcortical visual and auditory pathways, Lin et al. (*75*) proposed a two-module architecture for general spatiotemporal pattern recognition. A reservoir sequence encodes temporal patterns with different timescales into high-dimensional spatial representations, which are then interpreted by a decision-making module. This system has proven effective in tasks such as event-based gait recognition, outperforming deep learning methods under data-limited settings. Our approach here advances this concept by embedding domain-relevant inductive biases like direction selectivity and motion continuity–within the reservoir. These biases are introduced through a task-agnostic pretraining scheme inspired by the developmental processes of the animal visual system (*19, 20*). As a result, our pretrained recurrent network group capture more informative features relevant to real-world action recognition tasks, yielding stronger performance with greater efficiency.

#### Two-stream Convolutional Networks

These networks (*76*) represent a popular approach for video action recognition, using one CNN to process RGB frames and another to handle optical flow, with the outputs fused for prediction. While effective, this method relies heavily on external optical flow estimation, which is computationally expensive and often fails to capture all temporal cues (*77*). In contrast, our framework processes RGB videos directly via sparsely connected RNN modules (reservoirs), capturing rich temporal features without optical flow. This design minimizes training parameters, avoids full end-to-end training, and substantially reduces energy consumption, making it well-suited for few-shot learning and real-time applications on resource-limited devices.

#### Graph Neural Networks (GNNs)

GNNs have shown strong performance in skeleton-based action recognition by modeling human joints as nodes and their physical connections as edges (*78*). However, these methods face several limitations: 1) Reliance on keypoint detection, which is sensitive to occlusions and poor lighting; 2) Limited capacity for long-term temporal modeling, due to short processing windows; 3) Lack of interpretability, with learned weights often being opaque. Our framework addresses these limitations through the following: 1) Keypoint-free processing: By using RNNs to extract global motion patterns directly from raw frames, our method avoids reliance on explicit keypoint detection, increasing robustness to challenging video conditions; 2) Long-term temporal modeling: Pretrained RNN dynamics enable efficient capture of extended temporal dependencies critical for understanding complex actions; 3) Interpretability: The structured and modular design–grounded in biologically inspired mechanisms–makes the model transparent, facilitating insight into feature contributions and simplifying diagnostics.

#### Invariant feature learning

A central goal of domain adaptation is to learn invariant representations that generalize across distribution shifts. Data augmentation strategies (*36*) aim to approximate such invariance by synthetically perturbing inputs, but their effectiveness often depends on task-specific domain knowledge and access to large-scale labeled data. Contractive Auto-Encoders (CAEs) (*37*) promote local stability by penalizing the Jacobian of hidden activations with respect to inputs, thereby encouraging representations to lie on smooth, low-dimensional manifolds. Adversarial domain adaptation methods (*38–40, 79, 80*) align latent distributions across domains by training feature encoders to confuse a domain classifier via a gradient reversal mechanism. While effective in reducing domain-specific information, these approaches typically rely on large amounts of unlabeled target data and produce opaque features with limited interpretability. Causal representation learning (*41*) offers a principled alternative by identifying conditionally invariant mechanisms across environments. However, its practical deployment remains challenging due to the need for multiple domains, strong inductive biases, and assumptions about the underlying causal structure. Despite these advances, mainstream methods remain data-intensive, computationally heavy, and rely on black-box representations that are difficult to interpret or reuse. In contrast, our framework leverages evolutionarily conserved motion primitives and a simple curriculum-driven pretraining strategy to build explicit, reusable attractor-based features that align with the true physical invariance of the environment.

**Figure S1:**
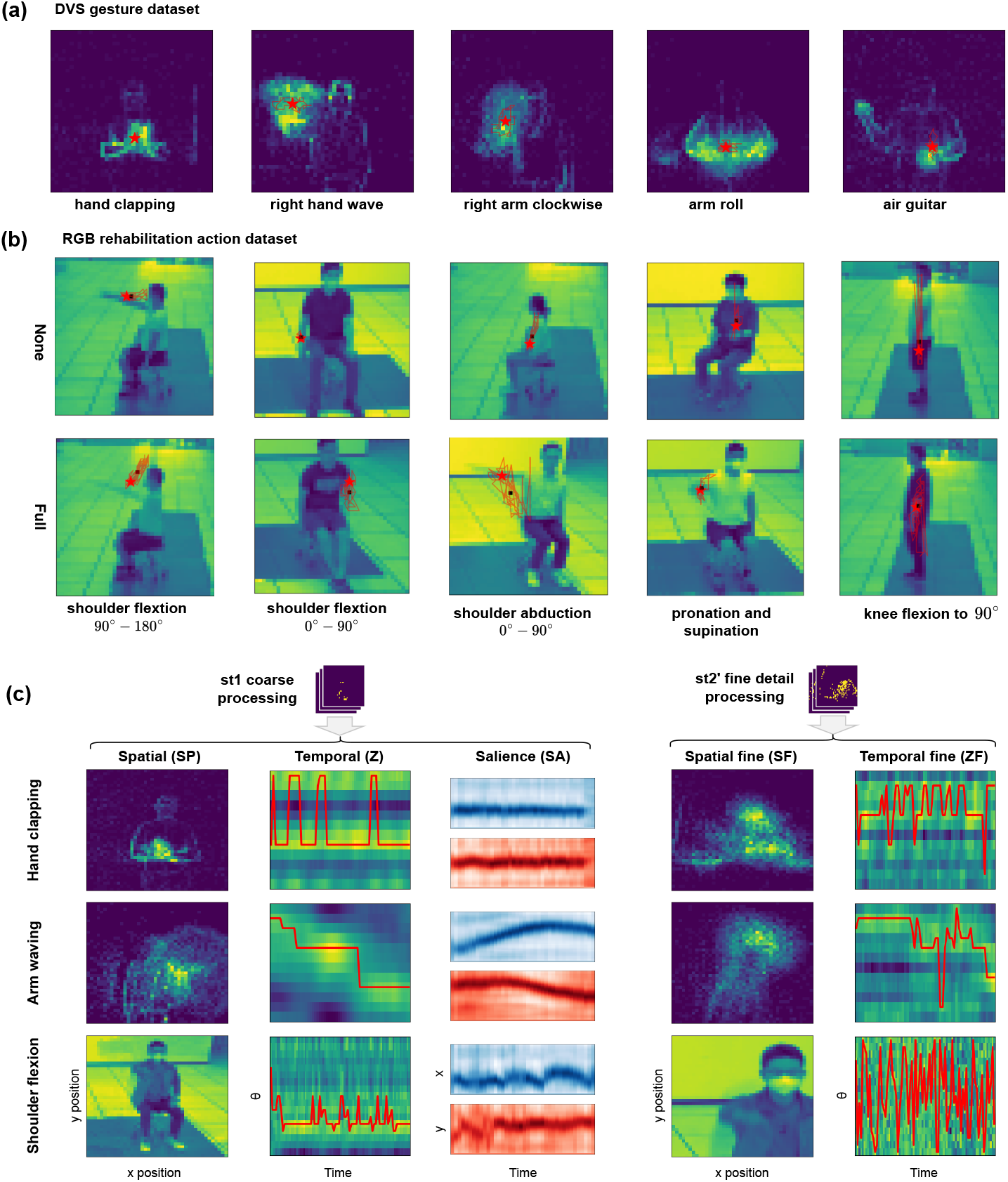
Dataset overview with salience focus position tracking. (**a**) DVS task for different categories with the focus trace provided by salience RNN(red line with the star indicating focus center across frames). (**b**) RGB rehabilitation action task for different categories with the focus trace provided by salience RNN(red line with the star indicating focus center across frames). For the RGB task, five major action categories are illustrated. (**c**) Visualizations of encoding features.

**Figure S2:**
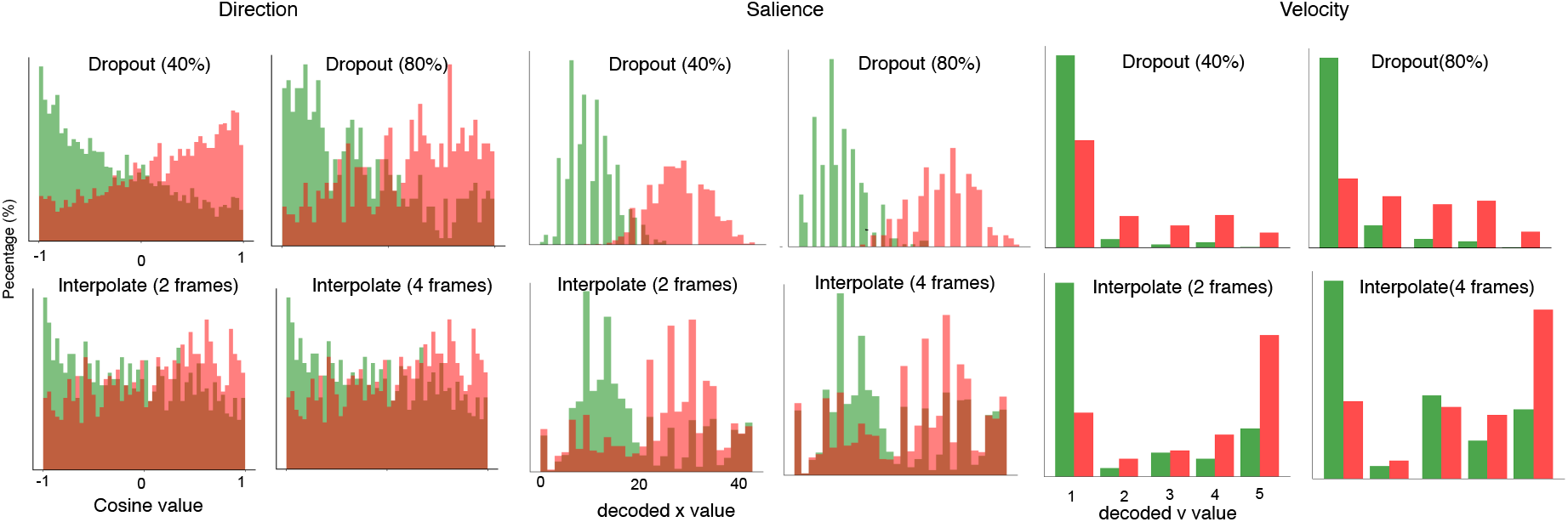
Measure of structure invariance across different noise condition. Dropout (40%, 80%) and interpolation (2 frames, 4 frames) for the Direction, Salience and Velocity RNNs.

## References and Notes

1. W. Xing, M. Li, M. Li, M. Han, Towards Robust and Secure Embodied AI: A Survey on Vulnerabilities and Attacks. arXiv preprint 2502.13175 (2025).

2. J. Liu, et al., Towards out-of-distribution generalization: A survey. arXiv preprint 2108.13624 (2021).

3. N. Roy, et al., From machine learning to robotics: Challenges and opportunities for embodied intelligence. arXiv preprint 2110.15245 (2021).

4. H. Jaeger, H. Haas, Harnessing nonlinearity: Predicting chaotic systems and saving energy in wireless communication. science 304 (5667), 78–80 (2004).

5. C. Barry, R. Hayman, N. Burgess, K. J. Jeffery, Experience-dependent rescaling of entorhinal grids. Nature neuroscience 10 (6), 682–684 (2007).

6. M. E. Thomason, et al., Cross-hemispheric functional connectivity in the human fetal brain. Science translational medicine 5 (173), 173ra24–173ra24 (2013).

7. M. Khona, I. R. Fiete, Attractor and integrator networks in the brain. Nature Reviews Neuroscience 23 (12), 744–766 (2022).

8. A. Nam, et al., Discrete, compositional, and symbolic representations through attractor dynamics. arXiv preprint 2310.01807 (2023).

9. M. N. Shadlen, W. T. Newsome, Neural basis of a perceptual decision in the parietal cortex (area LIP) of the rhesus monkey. Journal of neurophysiology 86 (4), 1916–1936 (2001).

10. Y. Burak, I. R. Fiete, Accurate path integration in continuous attractor network models of grid cells. PLoS computational biology 5 (2), e1000291 (2009).

11. M. W. Mathis, Adaptive Intelligence: leveraging insights from adaptive behavior in animals to build flexible AI systems. arXiv preprint 2411.15234 (2024).

12. G. Manjunath, P. Tino, H. Jaeger, Theory of input driven dynamical systems. ESANN 2012 proceedings (2012).

13. T. Nachstedt, C. Tetzlaff, Working memory requires a combination of transient and attractor-dominated dynamics to process unreliably timed inputs. Scientific reports 7 (1), 2473 (2017).

14. D. Sussillo, O. Barak, Opening the black box: low-dimensional dynamics in high-dimensional recurrent neural networks. Neural computation 25 (3), 626–649 (2013).

15. J. A. Gallego, M. G. Perich, L. E. Miller, S. A. Solla, Neural manifolds for the control of movement. Neuron 94 (5), 978–984 (2017).

16. G. Johansson, Visual perception of biological motion and a model for its analysis. Perception & psychophysics 14, 201–211 (1973).

17. K.-N. An, Kinematic analysis of human movement. Annals of biomedical engineering 12, 585–597 (1984).

18. R. J. Williams, D. Zipser, A learning algorithm for continually running fully recurrent neural networks. Neural computation 1 (2), 270–280 (1989).

19. Y. Li, D. Fitzpatrick, L. E. White, The development of direction selectivity in ferret visual cortex requires early visual experience. Nature neuroscience 9 (5), 676–681 (2006).

20. Y. Li, S. D. Van Hooser, M. Mazurek, L. E. White, D. Fitzpatrick, Experience with moving visual stimuli drives the early development of cortical direction selectivity. Nature 456 (7224), 952–956 (2008).

21. R. Pascanu, T. Mikolov, Y. Bengio, On the difficulty of training recurrent neural networks, in International conference on machine learning (Pmlr) (2013), pp. 1310–1318.

22. D. Ferster, K. D. Miller, Neural mechanisms of orientation selectivity in the visual cortex. Annual review of neuroscience 23 (1), 441–471 (2000).

23. M. A. Giese, T. Poggio, Neural mechanisms for the recognition of biological movements. Nature Reviews Neuroscience 4 (3), 179–192 (2003).

24. N. J. Priebe, Mechanisms of orientation selectivity in the primary visual cortex. Annual review of vision science 2 (1), 85–107 (2016).

25. S. Ji, W. Xu, M. Yang, K. Yu, 3D convolutional neural networks for human action recognition. IEEE transactions on pattern analysis and machine intelligence 35 (1), 221–231 (2012).

26. D. H. Hubel, T. N. Wiesel, Receptive fields, binocular interaction and functional architecture in the cat’s visual cortex. The Journal of physiology 160 (1), 106 (1962).

27. V. B. Mountcastle, The columnar organization of the neocortex. Brain: a journal of neurology 120 (4), 701–722 (1997).

28. N. J. Priebe, C. R. Cassanello, S. G. Lisberger, The neural representation of speed in macaque area MT/V5. Journal of Neuroscience 23 (13), 5650–5661 (2003).

29. J. D. Schall, On the role of frontal eye field in guiding attention and saccades. Vision research 44 (12), 1453–1467 (2004).

30. K. G. Thompson, K. L. Biscoe, T. R. Sato, Neuronal basis of covert spatial attention in the frontal eye field. Journal of Neuroscience 25 (41), 9479–9487 (2005).

31. J. I. Gold, M. N. Shadlen, The neural basis of decision making. Annu. Rev. Neurosci. 30 (1), 535–574 (2007).

32. M. O. Ernst, M. S. Banks, Humans integrate visual and haptic information in a statistically optimal fashion. Nature 415 (6870), 429–433 (2002).

33. A. Arnab, et al., Vivit: A video vision transformer, in Proceedings of the IEEE/CVF international conference on computer vision (2021), pp. 6836–6846.

34. A. Canatar, B. Bordelon, C. Pehlevan, Out-of-Distribution Generalization in Kernel Regression (2021).

35. S. Ben-David, et al., A theory of learning from different domains. Machine learning 79 (1), 151–175 (2010).

36. Y. Chen, et al., Understanding and Improving Feature Learning for Out-of-Distribution Generalization, in Thirty-seventh Conference on Neural Information Processing Systems (2023).

37. S. Rifai, P. Vincent, X. Muller, X. Glorot, Y. Bengio, Contractive auto-encoders: Explicit invariance during feature extraction, in Proceedings of the 28th international conference on international conference on machine learning (2011), pp. 833–840.

38. Y. Ganin, et al., Domain-adversarial training of neural networks. Journal of machine learning research 17 (59), 1–35 (2016).

39. J. Tobin, et al., Domain randomization for transferring deep neural networks from simulation to the real world, in 2017 IEEE/RSJ international conference on intelligent robots and systems (IROS) (IEEE) (2017), pp. 23–30.

40. Y. Shi, X. Ying, J. Yang, Deep unsupervised domain adaptation with time series sensor data: A survey. Sensors 22 (15), 5507 (2022).

41. M. Rojas-Carulla, B. Schölkopf, R. Turner, J. Peters, Invariant models for causal transfer learning. Journal of Machine Learning Research 19 (36), 1–34 (2018).

42. W. Maass, T. Natschläger, H. Markram, Real-time computing without stable states: A new framework for neural computation based on perturbations. Neural computation 14 (11), 2531– 2560 (2002).

43. A. Pascual-Leone, R. Hamilton, The metamodal organization of the brain. Progress in brain research 134, 427–445 (2001).

44. E. Marder, T. O’Leary, S. Shruti, Neuromodulation of circuits with variable parameters: single neurons and small circuits reveal principles of state-dependent and robust neuromodulation. Annual review of neuroscience 37 (1), 329–346 (2014).

45. R. J. Douglas, K. A. Martin, Neuronal circuits of the neocortex. Annu. Rev. Neurosci. 27 (1), 419–451 (2004).

46. S. Song, P.J. Sjöström, M. Reigl, S. Nelson, D. B. Chklovskii, Highly nonrandom features of synaptic connectivity in local cortical circuits. PLoS biology 3 (3), e68 (2005).

47. E. Rosch, M. Turbiaux, Classifications d’objets du monde réel: origines et représentations dans la cognition. Bulletin de psychologie 29 (325), 242–250 (1976).

48. B. M. Lake, T. D. Ullman, J. B. Tenenbaum, S. J. Gershman, Building machines that learn and think like people. Behavioral and brain sciences 40, e253 (2017).

49. A. O. Constantinescu, J. X. O’Reilly, T. E. Behrens, Organizing conceptual knowledge in humans with a gridlike code. Science 352 (6292), 1464–1468 (2016).

50. J. C. Whittington, et al., The Tolman-Eichenbaum machine: unifying space and relational memory through generalization in the hippocampal formation. Cell 183 (5), 1249–1263 (2020).

51. S. Wu, et al., Neural heterogeneity enhances reliable neural information processing: Local sensitivity and globally input-slaved transient dynamics. Science Advances 11 (14), eadr3903 (2025).

52. J. J. Hopfield, Neural networks and physical systems with emergent collective computational abilities. Proceedings of the national academy of sciences 79 (8), 2554–2558 (1982).

53. S.-i. Amari, Dynamics of pattern formation in lateral-inhibition type neural fields. Biological cybernetics 27 (2), 77–87 (1977).

54. Y. Bengio, A. Courville, P. Vincent, Representation learning: A review and new perspectives. IEEE transactions on pattern analysis and machine intelligence 35 (8), 1798–1828 (2013).

55. Y. LeCun, Y. Bengio, G. Hinton, Deep learning. nature 521 (7553), 436–444 (2015).

56. M. D. Zeiler, R. Fergus, Visualizing and understanding convolutional networks, in European conference on computer vision (Springer) (2014), pp. 818–833.

57. L. T. Oesch, M. C. Thomas, D. Sandberg, J. Couto, A. K. Churchland, Anterior cingulate cortex mixes retrospective cognitive signals and ongoing movement signatures during decisionmaking. bioRxiv pp. 2025–04 (2025).

58. R. B. Dekker, F. Otto, C. Summerfield, Curriculum learning for human compositional generalization. Proceedings of the National Academy of Sciences 119 (41), e2205582119 (2022).

59. A. Boopathy, et al., Breaking neural network scaling laws with modularity. arXiv preprint 2409.05780 (2024).

60. V. P. Pastore, P. Massobrio, A. Godjoski, S. Martinoia, Identification of excitatory-inhibitory links and network topology in large-scale neuronal assemblies from multi-electrode recordings. PLoS computational biology 14 (8), e1006381 (2018).

61. P. Werbos, Backpropagation through time: what it does and how to do it. Proceedings of the IEEE 78 (10), 1550–1560 (1990).

62. A. Amir, et al., A low power, fully event-based gesture recognition system, in Proceedings of the IEEE conference on computer vision and pattern recognition (2017), pp. 7243–7252.

63. D. J. Gladstone, C. J. Danells, S. E. Black, The Fugl-Meyer assessment of motor recovery after stroke: a critical review of its measurement properties. Neurorehabilitation and neural repair 16 (3), 232–240 (2002).

64. Z. Zhao, et al., The spike gating flow: A hierarchical structure-based spiking neural network for online gesture recognition. Frontiers in Neuroscience 16, 923587 (2022).

65. J. Redmon, S. Divvala, R. Girshick, A. Farhadi, You only look once: Unified, real-time object detection, in Proceedings of the IEEE conference on computer vision and pattern recognition (2016), pp. 779–788.

66. B. Böken, On the appropriateness of Platt scaling in classifier calibration. Information Systems 95, 101641 (2021).

67. H. Do, The organization of behavior. New York (1949).

68. F. Murtagh, P. Contreras, Algorithms for hierarchical clustering: an overview. Wiley Interdisciplinary Reviews: Data Mining and Knowledge Discovery 2 (1), 86–97 (2012).

69. D. Tran, L. Bourdev, R. Fergus, L. Torresani, M. Paluri, Learning spatiotemporal features with 3d convolutional networks, in Proceedings of the IEEE international conference on computer vision (2015), pp. 4489–4497.

70. B. Poole, S. Lahiri, M. Raghu, J. Sohl-Dickstein, S. Ganguli, Exponential expressivity in deep neural networks through transient chaos, in Advances in Neural Information Processing Systems 29, D. D. Lee, M. Sugiyama, U. V. Luxburg, I. Guyon, R. Garnett, Eds. (Curran Associates, Inc.), pp. 3360–3368 (2016).

71. W. Li, H. Wu, D. Yang, Powerful Encoding and Decoding Computation of Reservoir Computing, in International Conference on Intelligent Robotics and Applications (Springer) (2023), pp. 174–186.

72. D. Yang, T. Xu, F. Lin, Reservoir Computing in Rehabilitation Video Analyses, in 2023 IEEE International Conference on Cybernetics and Intelligent Systems (CIS) and IEEE Conference on Robotics, Automation and Mechatronics (RAM) (IEEE) (2023), pp. 210–214.

73. T. DeWolf, P. Jaworski, C. Eliasmith, Nengo and low-power AI hardware for robust, embedded neurorobotics. Frontiers in Neurorobotics 14, 568359 (2020).

74. J. Schmidhuber, S. Hochreiter, et al., Long short-term memory. Neural Comput 9 (8), 1735– 1780 (1997).

75. X. Lin, et al., A brain-inspired computational model for spatio-temporal information processing. Neural Networks 143, 74–87 (2021).

76. K. Simonyan, A. Zisserman, Two-stream convolutional networks for action recognition in videos. Advances in neural information processing systems 27 (2014).

77. Y. Zhang, et al., Gesture recognition based on deep deformable 3D convolutional neural networks. Pattern Recognition 107, 107416 (2020).

78. S. Yan, Y. Xiong, D. Lin, Spatial temporal graph convolutional networks for skeleton-based action recognition, in Proceedings of the AAAI conference on artificial intelligence, vol. 32 (2018).

79. Y. Ganin, V. Lempitsky, Unsupervised domain adaptation by backpropagation, in International conference on machine learning (PMLR) (2015), pp. 1180–1189.

80. X. B. Peng, M. Andrychowicz, W. Zaremba, P. Abbeel, Sim-to-real transfer of robotic control with dynamics randomization, in 2018 IEEE international conference on robotics and automation (ICRA) (IEEE) (2018), pp. 3803–3810.

